# Multiparametric Classification of Pure-tone Responses Distinguishes Neurons in Inferior Colliculus Subdivisions

**DOI:** 10.64898/2026.03.27.714825

**Authors:** Maryanna S. Owoc, Jongwon Lee, Adriana Johnson, Karl Kandler, Srivatsun Sadagopan

**Affiliations:** Department of Neurobiology, University of Pittsburgh School of Medicine, Pittsburgh, PA, USA; Center for Neuroscience, University of Pittsburgh, Pittsburgh, PA, USA; Medical Scientist Training Program, University of Pittsburgh/Carnegie Mellon University, Pittsburgh, PA, USA; Department of Bioengineering, University of Pittsburgh, Pittsburgh, PA, USA; Department of Communication Science and Disorders, University of Pittsburgh, Pittsburgh, PA, USA; Center for the Neural Basis of Cognition, Pittsburgh, PA, USA

## Abstract

The inferior colliculus (IC) integrates ascending auditory and descending multimodal inputs within distinct subdivisions, the central nucleus (CNIC) and cortex (CtxIC). Despite differences in connectivity, auditory responses in these subdivisions are similar, complicating localization during *in-vivo* recordings. Here, we tested whether recordings can be assigned to CNIC or CtxIC using only response properties in awake and anesthetized mice. We constructed frequency response areas (FRAs) from pure tone responses and extracted tuning and firing metrics. Individual FRA features could not reliably localize recordings. In contrast, a random forest classifier combining FRA-derived features accurately localized recordings to CNIC or CtxIC across states. These findings demonstrate that while IC subdivisions differ only subtly along individual response parameters, appropriate multiparametric approaches can enable robust classification. More broadly, our results illustrate how biologically meaningful distinctions may be revealed by combining weakly informative features, an approach that can be applied across diverse brain regions and modalities.

## INTRODUCTION

The inferior colliculus (IC) is a major hub for auditory processing in the midbrain, integrating ascending signals from the brainstem (Frisina et al., 1998) with descending cortical projections (Saldaña et al., 1996; Winer, 2006), commissural projections (Chandrasekaran et al., 2013; Ito and Oliver, 2014; Malmierca et al., 2003, 1995; Rees and Orton, 2019; Saldaña and Merchán, 1992), and intrinsic circuitry (Ito and Oliver, 2014; Malmierca et al., 1995; Miller et al., 2005; Oliver et al., 1991; Saldaña and Merchán, 1992; Sturm et al., 2014, 2017; Wallace et al., 2012). Anatomically, the IC is organized into a ‘core’ central nucleus (CNIC) which Is surrounded by a ‘shell’, or cortex (CtxIC). The CtxIC can further be subdivided into the dorsal and lateral cortices. The CNIC receives predominantly ascending inputs from lower brainstem nuclei (Beyerl, 1978; Coleman and Clerici, 1987; Shneiderman et al., 1988; Sitek et al., 2022) and accounts for a substantial portion of commissural projections (Rees and Orton, 2019). The lateral CtxIC integrates multimodal information, including somatosensory and visual inputs, and descending auditory signals (Coleman and Clerici, 1987; Faye-Lund, 1985; Lesicko et al., 2016; Li and Mizuno, 1997; Wise and Jones, 1977). The dorsal cortex receives inputs from the auditory cortex, cochlear nucleus, and the dorsal nucleus of the lateral lemniscus (Coleman and Clerici, 1987; Druga et al., 1997; Druga and Syka, 1984; Faye-Lund, 1985; Sitek et al., 2022). This differential connectivity of the subregions suggests that CNIC and CtxIC should support distinct computations, and as a consequence, exhibit distinguishable response properties.

But paradoxically, single-neuron electrophysiological responses recorded *in-vivo* from these subdivisions are reported to be quite similar. Many studies have focused primarily on the CNIC (Egorova et al., 2006, 2001, 2020; Lee et al., 2019) or pooled data across IC subdivisions without precise localization (Galazyuk et al., 2017; Ono and Oliver, 2014; Portfors and Felix, 2005; Tan et al., 2007; Walton et al., 1997), limiting the ability to determine responses differences across subdivisions. A few studies across species (cats: Aitkin et al., 1994, 1981, 1978; guinea pigs: Syka et al., 2000; mice: Barnstedt et al., 2015; Wong and Borst, 2019) have reported some differences such as broader tuning in CtxIC or different extents of monotonic rate-level functions. However, the distributions of these parameters typically show substantial overlap across subdivisions, and whether these differences would support classifying recording location is unclear. Furthermore, much of the prior work has been performed under anesthesia using different agents, potentially masking subdivision-specific differences or introducing anesthetic-dependent biases. Together, these factors have complicated the reliable assignment of recording location to IC subdivisions based solely on response properties.

Accurate online localization of recording locations to IC subdivisions is critical for circuit-dissection experiments. Examples of such experiments include injections of viral vectors, pharmacological agents, or precursor neurons (Owoc et al., 2022), lesioning or inactivation of specific subdivisions, and microstimulation of functionally characterized sites. In many species, especially those in which the IC is deep and not reliably accessible using stereotactic coordinates such as non-human primates (Rocchi and Ramachandran, 2018; Slee and Young, 2011; Wang et al., 2022), physiological response properties are essential for guiding these recordings and circuit manipulations. A robust, generalizable method for classifying CNIC versus CtxIC location from neural response features would enable experiments across preparations and states and enhance the precision of *in-vivo* studies of auditory circuit function.

Here, we address this need by asking two questions: (1) Can neurons be reliably assigned to CNIC or CtxIC using only their auditory response properties?, and (2) Is such classification robust to anesthesia? To answer these questions, we recorded from histologically verified CNIC and CtxIC neurons in awake and isoflurane-anesthetized mice and collected responses to pure tones and dynamic random chord (DRC) stimuli. From pure-tone responses, we constructed frequency response areas (FRAs) and extracted tuning (e.g., threshold, bandwidth) and firing metrics (e.g., spontaneous and driven rates). We also estimated spectrotemporal receptive fields (STRFs) from responses to DRC stimuli to assess spectrotemporal differences. Consistent with prior observations, individual FRA-derived features and STRF parameters were largely overlapping between CNIC and CtxIC neurons along single dimensions. In contrast, machine-learning approaches, particularly those using non-linear combinations of FRA features such as random forest (RF) classifiers, robustly localized neurons to CNIC or CtxIC in both awake and anesthetized recordings. Anesthesia increased response reliability, differentially affected thresholds across subdivisions, and increased classifier performance. But high performance remained after excluding threshold as a feature, indicating that the multiparametric benefit was not driven by a single anesthetic-sensitive parameter.

Together, these results reveal that CNIC and CtxIC differ only subtly along single response features and that appropriately combining these features enables reliable and accurate localization based on response properties. More broadly, our study highlights the utility of machine learning tools for online targeting of other brain structures (cortical areas, thalamic nuclei) and cell-type inference based on response parameters, providing a framework to increase the precision of *in-vivo* circuit studies.

## RESULTS

*In-vivo* extracellular recordings were obtained from the ICs of adult (6 – 13 weeks) awake and isoflurane-anesthetized CBA/CaJ mice (Jax 000654) using 64-channel silicon microelectrode arrays (Du et al., 2011). For awake experiments, recordings were performed in 17 head-fixed mice (n = 17 ICs, 28 tracks; Fig. 1A), of which 14 tracks only contained CNIC units, 10 tracks only contained CtxIC units, and 4 tracks contained both CNIC and CtxIC units. For anesthetized experiments, recordings were performed in 5 isoflurane-anesthetized mice (n = 7 ICs), of which 4 tracks were confirmed to be in the CNIC and 3 in the CtxIC. The ground-truth recording location was determined histologically by coating the probe with a fluorescent dye (DiI or DiO) and delineating the CNIC using cytochrome oxidase staining (see Methods; Fig. 1B,C). In cases where a track spanned both subdivisions, we determined the depth at which the probe crossed the CtxIC/CNIC boundary (Fig. 1B) and assigned single units to IC subdivisions accordingly. In several CNIC tracks, we observed an expected dorsoventral tonotopic gradient in multiunit responses, further supporting localization to CNIC (Fig. 1D). From these recordings, we used established spike-sorting software (Kilosort4 and JRCLUST) and stringent criteria (SNR > 4, <10% refractory period violations, stable firing rate across recording duration) to isolate single-unit clusters (Fig. 1E). For the FRA-based analyses, we isolated 87 CNIC and 65 CtxIC tone-responsive units in the awake mice, and 135 CNIC and 125 CtxIC tone-responsive units in the anesthetized mice, with best frequencies (BFs) ranging from 4 to 32 kHz (Fig. 1D) and spanning a wide range of thresholds and best levels (BLs).

**Figure 1:**
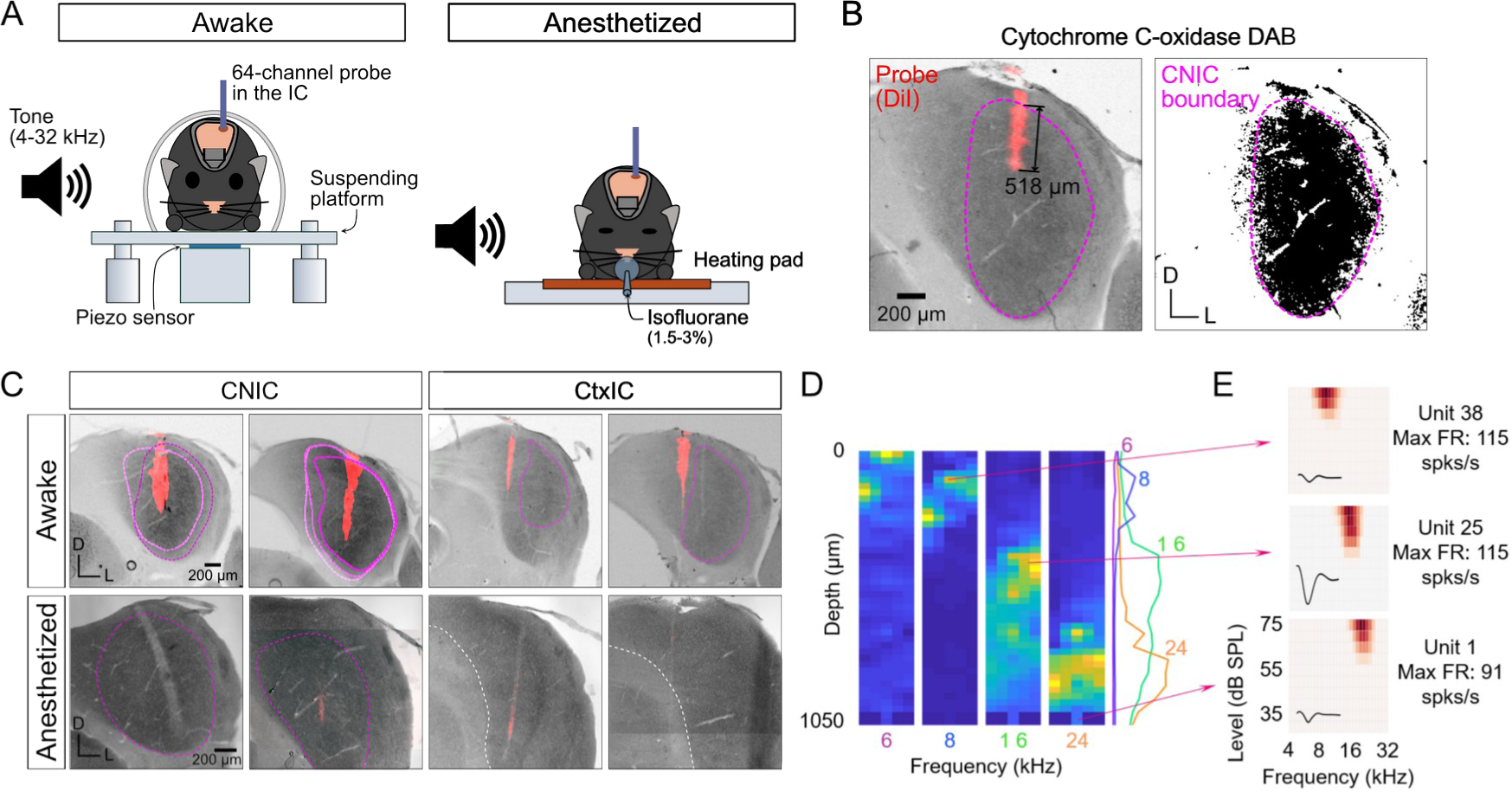
Electrophysiological recordings and probe track localization. **(A)** Schematic of recording setup for awake and anesthetized recordings. **(B)** Histological image showing delineation of CNIC and CtxIC in an IC section (magenta line) using cytochrome oxidase staining. Thresholded image on right. Red staining is the fluorescent dye (DiI) deposited by the electrode track. **(C)** IC sections showing electrode tracks localized to CNIC and CtxIC. **(D)** Heatmap showing multiunit activity in response to tones of different frequencies. Many CNIC tracks exhibited an expected dorsoventral tonotopic gradient. **(E)** Example single-units and their FRAs from different depths of a CNIC track.

### CNIC and CtxIC units exhibit similar spontaneous and driven firing profiles for each state

We began by examining overall spontaneous and tone-driven response properties of CNIC and CtxIC units. For readability, the medians and 95% confidence intervals of all parameters described below are summarized in **Table 1**. Example spike rasters of CNIC neurons in the awake and anesthetized state are plotted in Fig. 2A and 2G respectively. We found no significant difference between the spontaneous firing rates of CNIC and CtxIC neurons in awake or anesthetized mice. While spontaneous rates in both CNIC and CtxIC were much lower in the anesthetized state compared to the awake state, the lack of a difference between the two subdivisions is in contrast to a previous study in urethane- and ketamine-anesthetized guinea pigs (Syka et al., 2000). The maximal tone-driven firing rates, measured at BF and BL, were similarly higher in the awake state for both CNIC and CtxIC neurons compared to the anesthetized state, with CNIC neurons exhibiting significantly higher max. driven rates across both states (Fig. 2D,J). We found no differences in the onset latencies of CNIC and CtxIC neurons in the awake and anesthetized states, but latencies in the anesthetized state were lower than in the awake state (Fig. 2E,K). We attribute this difference to our method of estimating latency (see Methods), which is sensitive to ongoing spontaneous activity that is higher in the awake state. We then analyzed the temporal profile of sound-driven responses by computing an onset/offset ratio (see Methods), derived from the onset (0 – 50 ms window after tone onset) and offset (0 – 50 ms window after tone offset) firing rates at BF and BL for each neuron. While this index was also not discriminative between CNIC and CtxIC neurons in both awake and anesthetized states, we observed a striking difference between the awake and anesthetized states (Fig. 2F,L). In awake mice, we encountered neurons with a range of onset and offset response magnitudes (Fig. 2F), whereas we encountered little to no offset responses in isoflurane-anesthetized mice (Fig. 2L). In summary, only the maximal driven rate was significantly different (at p<0.01) between CNIC and CtxIC neurons in the awake state, and response parameters did not reliably distinguish between CNIC and CtxIC recording locations in either state.

**Table 1:**
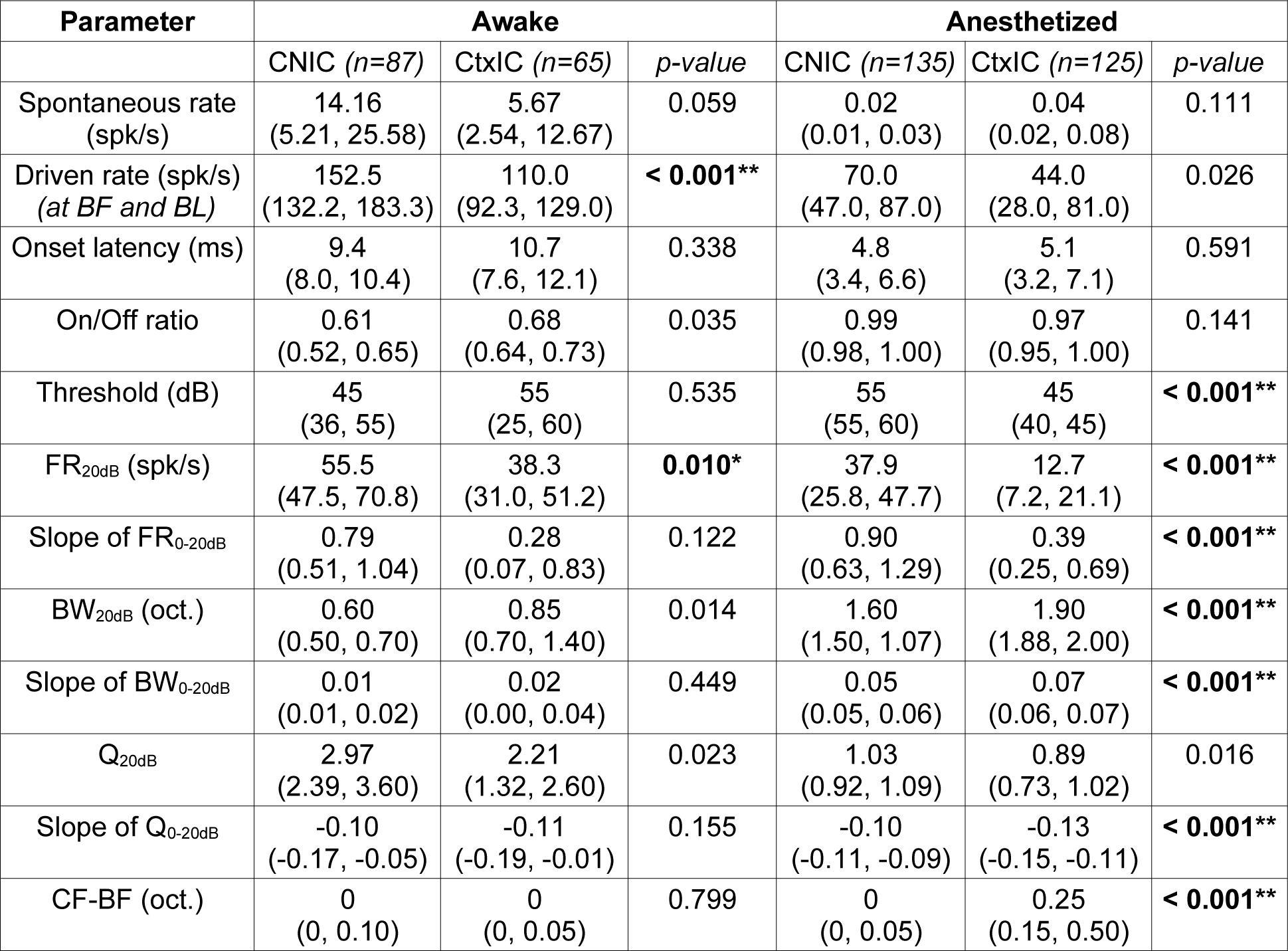
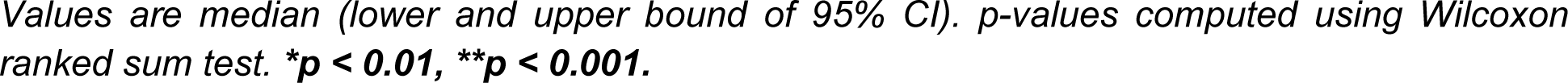
Summary of firing rate and FRA response properties.

| Parameter | Awake |  |  | Anesthetized |  |  |
| --- | --- | --- | --- | --- | --- | --- |
|  | CNIC (n=87) | CtxIC (n=65) | p-value | CNIC (n=135) | CtxIC (n=125) | p-value |
| Spontaneous rate (spk/s) | 14.16<br>(5.21, 25.58) | 5.67<br>(2.54, 12.67) | 0.059 | 0.02<br>(0.01, 0.03) | 0.04<br>(0.02, 0.08) | 0.111 |
| Driven rate (spk/s)<br>(at BF and BL) | 152.5<br>(132.2, 183.3) | 110.0<br>(92.3, 129.0) | **< 0.001\*\*** | 70.0<br>(47.0, 87.0) | 44.0<br>(28.0, 81.0) | 0.026 |
| Onset latency (ms) | 9.4<br>(8.0, 10.4) | 10.7<br>(7.6, 12.1) | 0.338 | 4.8<br>(3.4, 6.6) | 5.1<br>(3.2, 7.1) | 0.591 |
| On/Off ratio | 0.61<br>(0.52, 0.65) | 0.68<br>(0.64, 0.73) | 0.035 | 0.99<br>(0.98, 1.00) | 0.97<br>(0.95, 1.00) | 0.141 |
| Threshold (dB) | 45<br>(36, 55) | 55<br>(25, 60) | 0.535 | 55<br>(55, 60) | 45<br>(40, 45) | **< 0.001\*\*** |
| FR<sub>20dB</sub> (spk/s) | 55.5<br>(47.5, 70.8) | 38.3<br>(31.0, 51.2) | **0.010\*** | 37.9<br>(25.8, 47.7) | 12.7<br>(7.2, 21.1) | **< 0.001\*\*** |
| Slope of FR<sub>0-20dB</sub> | 0.79<br>(0.51, 1.04) | 0.28<br>(0.07, 0.83) | 0.122 | 0.90<br>(0.63, 1.29) | 0.39<br>(0.25, 0.69) | **< 0.001\*\*** |
| BW<sub>20dB</sub> (oct.) | 0.60<br>(0.50, 0.70) | 0.85<br>(0.70, 1.40) | 0.014 | 1.60<br>(1.50, 1.07) | 1.90<br>(1.88, 2.00) | **< 0.001\*\*** |
| Slope of BW<sub>0-20dB</sub> | 0.01<br>(0.01, 0.02) | 0.02<br>(0.00, 0.04) | 0.449 | 0.05<br>(0.05, 0.06) | 0.07<br>(0.06, 0.07) | **< 0.001\*\*** |
| Q<sub>20dB</sub> | 2.97<br>(2.39, 3.60) | 2.21<br>(1.32, 2.60) | 0.023 | 1.03<br>(0.92, 1.09) | 0.89<br>(0.73, 1.02) | 0.016 |
| Slope of Q<sub>0-20dB</sub> | -0.10<br>(-0.17, -0.05) | -0.11<br>(-0.19, -0.01) | 0.155 | -0.10<br>(-0.11, -0.09) | -0.13<br>(-0.15, -0.11) | **< 0.001\*\*** |
| CF-BF (oct.) | 0<br>(0, 0.10) | 0<br>(0, 0.05) | 0.799 | 0<br>(0, 0.05) | 0.25<br>(0.15, 0.50) | **< 0.001\*\*** |

Values are median (lower and upper bound of 95% CI). *p*-values computed using Wilcoxon ranked sum test. \**p* < 0.01, \*\**p* < 0.001.

**Figure 2:**
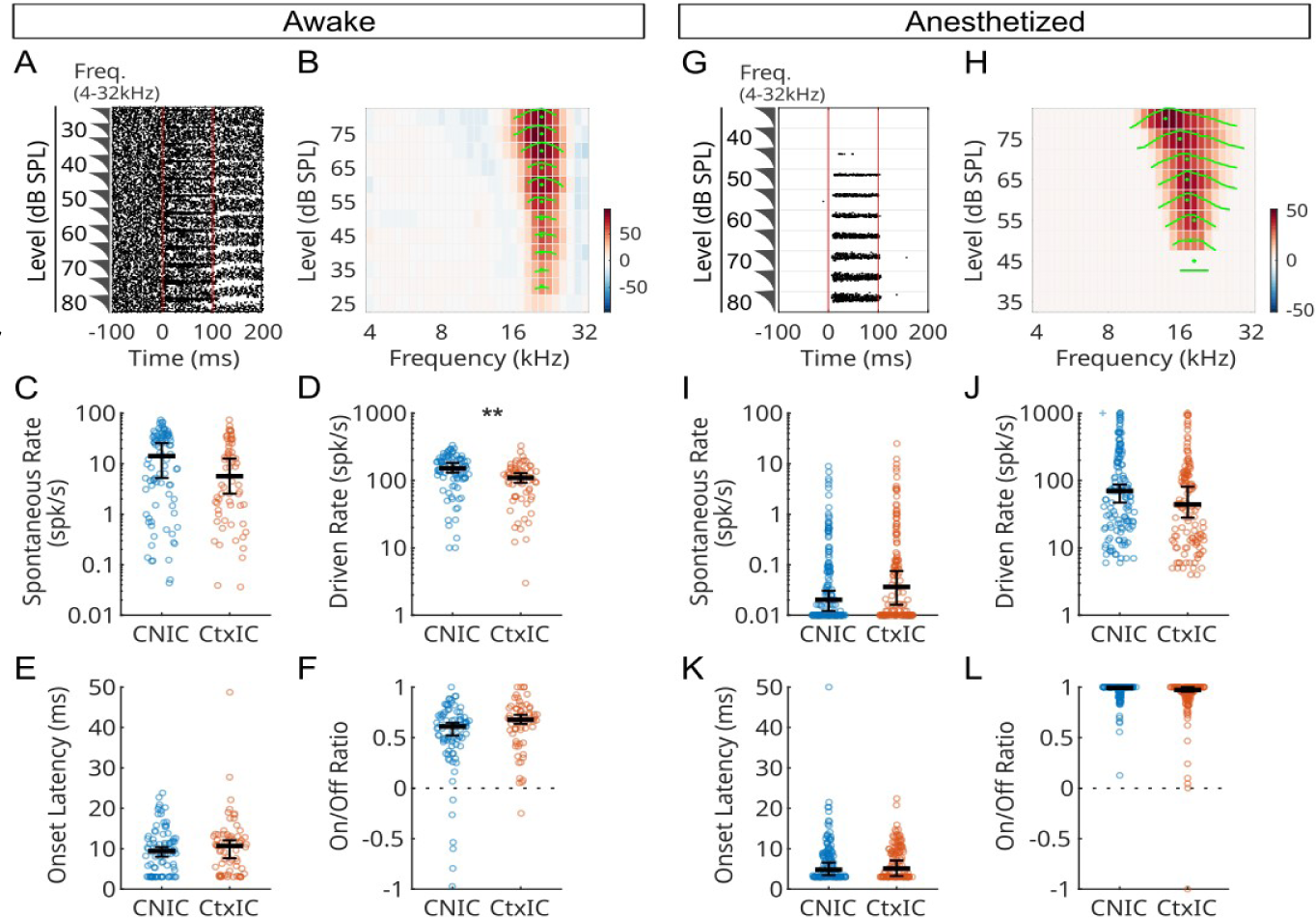
Response properties across IC subdivisions and states. **Awake: (A)** Representative spike raster of a CNIC neuron showing high spontaneous rate, sustained stimulus-driven response, and post-stimulus activity. Red lines indicate stimulus duration. Note that frequency and sound level are interleaved on the y-axis, with the sawtooth schematic representing stimulus frequency. **(B)** FRA computed from the raster in A. Colormap corresponds to firing rate computed over the entire stimulus duration. **(C)** Mean spontaneous rate of CNIC (blue) and CtxIC (red) neurons. **(D)** Mean onset latency, estimated as the knee point of a piecewise linear function. **(E)** Maximum driven rates of neurons in CNIC and CtxIC. **(F)** Offset index, computed as in Eqn. X, showing a range of temporal response profiles in both CNIC and CtxIC. **Anesthetized: (G)** Representative spike raster of a CNIC neuron showing very low spontaneous rate, sustained stimulus-driven response, and no post-stimulus activity. **(H – L)** Same as (B – F) but in anesthetized mice. (*+: Data point outside range; **: p<0.001, see Table 1 for statistical details*).

### CNIC and CtxIC units exhibit similar FRAs for each state

We then examined the FRAs of neurons across the two IC subunits. In both IC subunits and in both states, we sampled neurons with best frequencies ranging from 4 to 32 kHz (Fig. 3A,M). The FRAs of CNIC and CtxIC units were largely similar in shape for each state (Fig. 2B,H; CNIC FRAs shown), but in contrast to data from decerebrate cats (Ramachandran et al.,1999), “V” shaped FRAs were the most prevalent and more likely to be encountered in CNIC in the anesthetized state (CNIC: 91%, CtxIC 62%) than the awake state (CNIC: 59%, CtxIC: 25%). We derived numerous parameters to characterize the FRAs of neurons across IC subunits and states (Fig. 3). For readability, the medians and 95% confidence intervals of these parameters are summarized in **Table 1**. In the awake state, the thresholds of neurons were similar across both CNIC and CtxIC (Fig. 3B), but under anesthesia, thresholds in both IC subdivisions were elevated and CNIC neurons showed higher thresholds than CtxIC neurons (Fig. 3N), likely reflecting the differential effects of isoflurane anesthesia across IC subdivisions. Close to threshold (20 dB above threshold), CNIC units exhibited narrow frequency tuning (Fig. 3H) which was subtly narrower than CtxIC tuning bandwidths (p = 0.014). At higher sound levels, however, we did not observe systematic differences in CNIC and CtxIC bandwidths in the awake state (Fig. 3G). Under anesthesia, CNIC and CtxIC bandwidths were both consistently broader than in the awake state (Fig. 3T). It should be noted, however, that the threshold was also higher under anesthesia (making the same relative sound levels be of higher absolute amplitude). Consistent with earlier studies (Aitkin et al., 1975; Syka et al., 2000), under anesthesia, CtxIC neurons displayed higher bandwidths than CNIC neurons (Fig. 3S).

**Figure 3:**
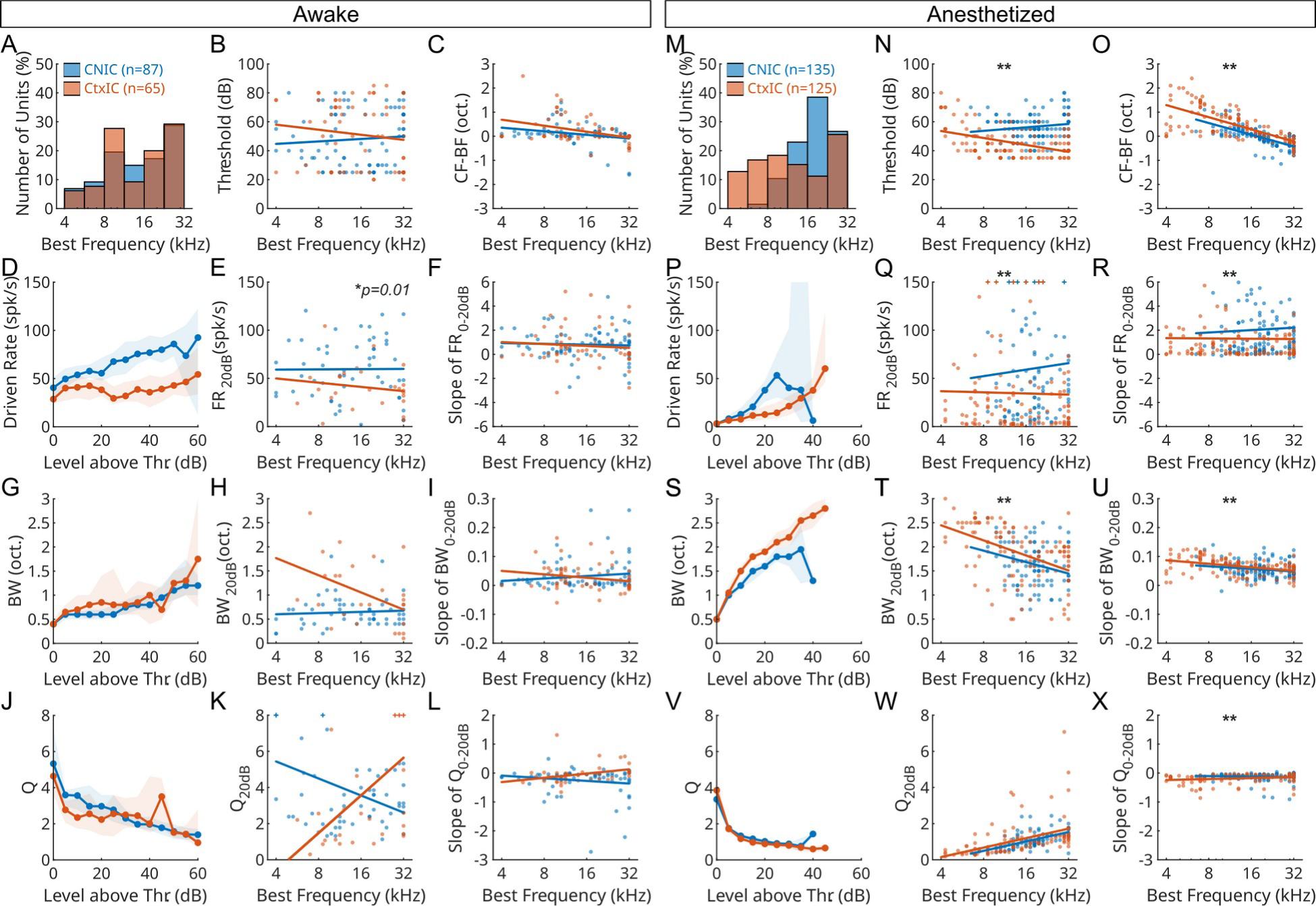
FRA parameters across IC subdivisions and states. **Awake: (A)** Distribution of the best frequencies of recorded CNIC (blue) and CtxIC (red) units. **(B)** Response thresholds of individual CNIC and CtxIC units (dots) and linear fits as a function of best frequency. **(C)** Symmetry of FRAs, characterized as the difference between the characteristic and best frequencies. **(D – F)** Characterization of driven rates of CNIC and CtxIC units as a function of suprathreshold sound level and at 20 dB above threshold. **(G – I)** Characterization of the bandwidths of CNIC and CtxIC units as a function of suprathreshold sound level and at 20 dB above threshold. **(J – L)** Characterization of the quality factors of CNIC and CtxIC units as a function of suprathreshold sound level and at 20 dB above threshold. **Anesthetized: (M – X)** Same as (A – L) but under isoflurane anesthesia. (*+: Data point outside plot range; *: p≤0.01, **: p<0.001, see Table 1 for statistical details*).

We also characterized tuning symmetry as the difference between the BF and the characteristic frequency (CF; the BF at threshold). In both IC subdivisions and across states, FRAs exhibited a strong asymmetry (CF-BF≥0) that indicated a low-frequency tail (Fig. 3C,O). This asymmetry was more pronounced under anesthesia. Most FRAs showed monotonically increasing rate-level functions (Fig. 3D,P), and while CNIC neurons tended to exhibit higher median firing rates at 20 dB above threshold than CtxIC neurons (Fig. 3E,Q), this difference was statistically significant only in the anesthetized state. The slopes of firing rate increase with sound level were indistinguishable between IC subdivisions in awake animals, but CNIC neurons showed significantly steeper slopes under anesthesia (Fig. 3F,R). To characterize how tuning bandwidth (BW) increased with increasing stimulus level, we plotted the BW in octaves at each level relative to threshold and measured the bandwidth at 20 dB above threshold and the rate of increase of bandwidth between the threshold and 20 dB above threshold. For these parameters as well, we observed no significant differences between IC subdivisions in awake animals. In anesthetized animals, BWs of CtxIC neurons were broader than those of CNIC neurons at louder sound levels (Fig. 3T). Consistent with these observations, the Q value and the rate of growth of Q with sound level were also not different between IC subdivisions in awake animals. These results are summarized in Table 1. Overall, this exhaustive characterization demonstrates that in the awake state, FRA parameters of CNIC and CtxIC neurons are largely similar and do not support reliable localization of recordings based on receptive field parameters alone. Anesthesia, likely through differential mechanisms of action across IC subdivisions, introduced statistically significant differences between CNIC and CtxIC responses along many of these FRA parameters. But as we will demonstrate later in this manuscript (Fig. 6), effect sizes along individual parameters were not sufficient to support reliable localization of recordings to CNIC and CtxIC in anesthetized recordings as well.

### CNIC and CtxIC units exhibit similar STRFs for each state

Presenting single pure-tones does not engage known mechanisms such as side-band suppression, and firing rate analyses based on relatively static and long-duration stimuli do not capture the temporal dynamics of neural responses. To address these limitations, we used dynamic random chord (DRC) stimuli (Fig. 4A) to evoke responses (Fig. 4B), and fit linear-nonlinear models to the responses using the Neural Encoding Model System (NEMS) (Pennington and David, 2020; Thorson et al., 2015) to determine spectrotemporal receptive fields (STRFs). For each recorded unit, the encoding model estimated a set of 360 linear weights (36 frequencies x 10 time bins, taken to be the STRF, Fig. 4C,E) and the parameters (n=5) of a level-shifted double-exponential static nonlinearity. Neural responses to the DRC stimulus were predicted using this model with nested cross-validation (see Methods), and the correlation coefficient between the predicted PSTH and the actual PSTH (r-test) was computed (see example in Fig. 4B). STRFs with low r-test values (r-test<0.3) were excluded from further analysis (dashed line in Fig. 4D,F). The median r-test of the included STRFs was not different between CNIC and CtxIC across states (Fig. 4D,F) indicating that there were no systematic differences in the convergence of STRF estimation procedures between the two regions. Median r-test values were lower for the awake state, potentially reflecting the influence of non-auditory modulations of neural activity (Yang et al., 2020; Saderi et al., 2021; Quass et al., 2024), which our encoding model was not designed to capture.

**Figure 4.**
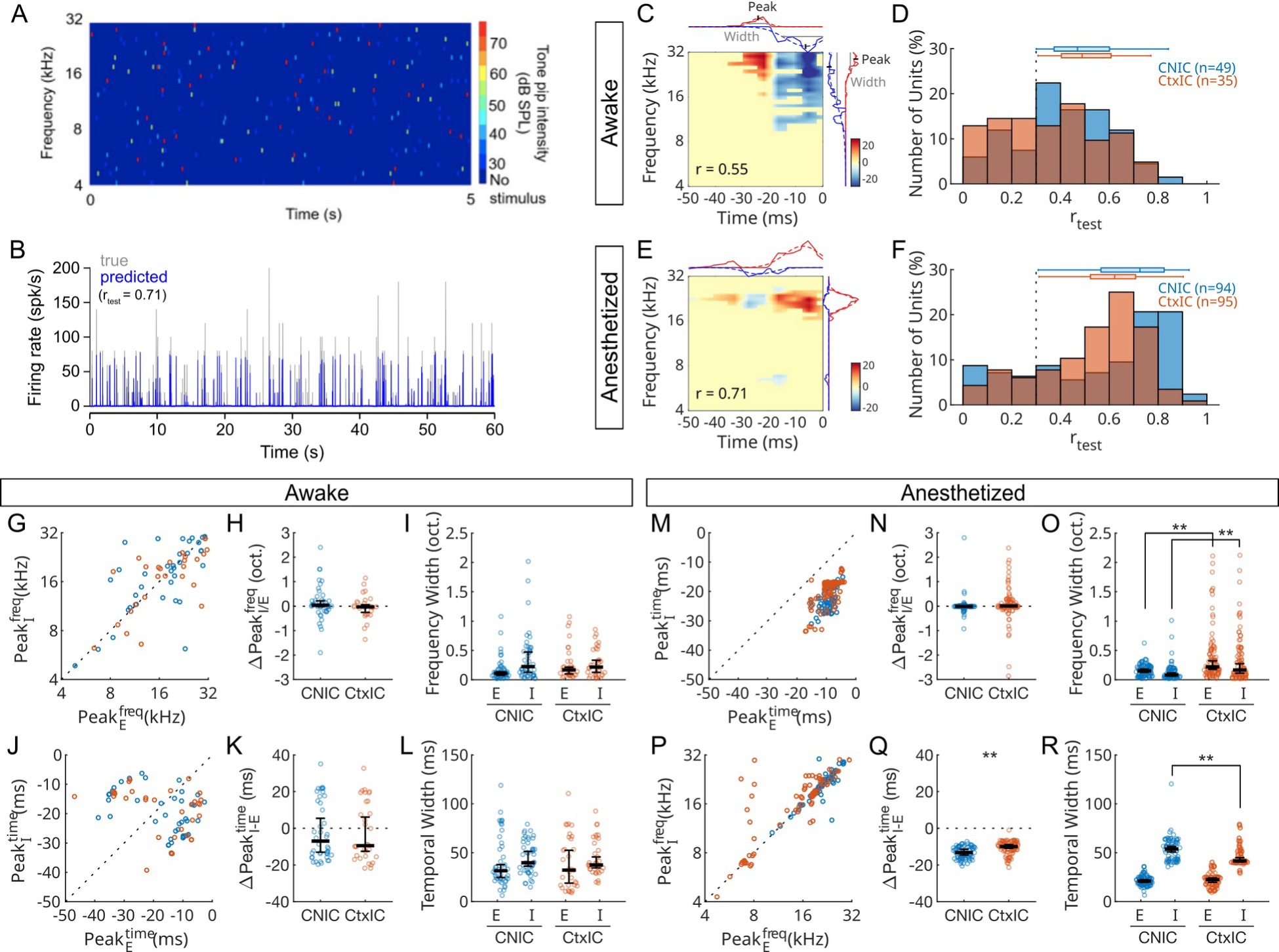
Comparison of STRF parameters of CNIC and CtxIC units in the awake and anesthetized states. **(A)** A 5 second segment of the STRF stimulus, a dynamic random chord (DRC). Colors correspond to tone pip intensity (right margin: color map scale, dark blue – no stimulus, red – 75 dB SPL). **(B)** Actual response PSTH (blue) and the predicted response PSTH (orange) using the STRF shown in E. The correlation coefficient between the predicted and the actual response (r-test) was 0.7. **(C, E)** Representative STRFs from awake and anesthetized recordings showing excitatory (red) and inhibitory (blue) subunits. The frequency (right) and temporal (top) response profiles for both the excitatory (red) and inhibitory (blue) response are shown along the margins. The position (mean) and width (standard deviation) of the response profiles were derived from Gaussian fits. **(D, F)** Distribution of the correlation values (r-test) between the predicted and actual PSTH. Single units with a r-test less than 0.3 (dashed line) were excluded from further analysis. The median and interquartile range for each distribution, after excluding r<0.3, are plotted above the distributions. CNIC is blue, CtxIC is red. **Awake: (G – H)** Summary of STRF parameters in the awake state for CNIC (blue) and CtxIC (red) units. **(G)** Relative position of the excitatory and inhibitory peaks in the frequency axis. **(H)** Distributions of the frequency differences between excitatory and inhibitory frequency peaks. **(I)** Frequency bandwidths of the excitatory and inhibitory STRF subunits. **(J)** Relative position of the excitatory and inhibitory peaks in the time axis. **(K)** Distributions of the frequency differences between excitatory and inhibitory peaks in the time axis. **(L)** Temporal durations of the excitatory and inhibitory STRF subunits. **Anesthetized: (M – R)** Same as (G – H) but for STRFs in the anesthetized state. (*\*\*: p<0.001, see Table 2 for statistical details*).

**Table 2:** Summary of STRF properties.

| Parameter | Awake |  |  | Anesthetized |  |  |
| --- | --- | --- | --- | --- | --- | --- |
|  | CNIC (n=49) | CtxIC (n=35) | *p*-value | CNIC (n=94) | CtxIC (n=95) | *p*-value |
| E<sup>freq</sup> width (oct.) | 0.11<br>(0.07, 0.14) | 0.17<br>(0.10, 0.21) | 0.030 | 0.15<br>(0.13, 0.18) | 0.22<br>(0.18, 0.32) | < 0.001** |
| I<sup>freq</sup> width (oct.) | 0.22<br>(0.13, 0.47) | 0.22<br>(0.13, 0.33) | 0.684 | 0.08<br>(0.06, 0.12) | 0.17<br>(0.11, 0.27) | < 0.001** |
| ΔPeak<sub>I/E</sub><sup>freq</sup> (oct.) | 0.04<br>(-0.02, 0.22) | -0.03<br>(-0.25, 0.06) | 0.075 | -0.01<br>(-0.01, 0) | 0.01<br>(-0.02, 0.04) | 0.140 |
| E<sup>time</sup> width (ms) | 31.6<br>(24.5, 37.8) | 32.2<br>(18.9, 52.5) | 0.418 | 20.9<br>(19.7, 22.4) | 22.4<br>(19.7, 24.1) | 0.717 |
| I<sup>time</sup> width (ms) | 39.8<br>(36.1, 51.3) | 37.4<br>(34.5, 45.7) | 0.890 | 53.8<br>(50.8, 56.4) | 41.6<br>(40.6, 44.9) | < 0.001** |
| ΔPeak<sub>I/E</sub><sup>time</sup> (ms) | -6.9<br>(-13.0, 5.4) | -9.4<br>(-12.5, 6.1) | 0.655 | -13.2<br>(-14.6, -11.4) | -9.9<br>(-10.9, -8.9) | < 0.001** |

Values are median (lower and upper bound of 95% CI). *p*-values computed using Wilcoxon ranked sum test. \*\*: *p* < 0.001.

From each STRF, by fitting Gaussians to the positive and negative STRF weights averaged separately along the frequency and time axes, we extracted four ‘tuning curves’ corresponding to the excitatory (E) and inhibitory (I) spectral and temporal response profiles (red (E) and blue (I) curves along the margins of the STRFs in Fig. 4C,E). Eight parameters were extracted to quantify these tuning curves – the peak E and I position in frequency and time (corresponding to the means of the Gaussian fits), and the width of E and I in frequency and time (corresponding to the full width at half maximum of the Gaussian fits). Response profiles with peak positions that fell out of the range of STRF frequency and time extents were excluded from analysis.

Surprisingly, rather than lateral inhibition peaks that straddled an excitatory peak, most units in both IC subdivisions and in both states exhibited co-tuned E and I frequency tuning curves (Fig. 4G,M). The E and I subunits were coincident in frequency but were temporally displaced, i.e., with E or I leading (Fig. 4J,P). E and I bandwidths were significantly different between CNIC and CtxIC units in the anesthetized state, but not in the awake state. Consistent with the FRA analysis described above, E frequency tuning widths were slightly narrower in CNIC units compared to CtxIC units across both states. Interestingly, I frequency tuning widths were significantly smaller in the CNIC compared to the CtxIC only in the anesthetized state, hinting at differential effects of isoflurane on excitatory and inhibitory circuits. Because we used a spectrotemporally varying stimulus (DRC), these data also allowed us to characterize the time course of E and I responses. In the awake state, consistent with the response profile analysis described in Fig. 2F, we found units in both CNIC and CtxIC with E-leading as well as I-leading STRF profiles (Fig. 4J), resulting in broad distributions of the differences in the peak times of the E and I subunits (Fig. 4K). In stark contrast, we exclusively observed onset type responses in the awake state (Fig. 4P,Q). While statistically significant differences emerged in the temporal extent of I subunits between CNIC and CtxIC neurons in the anesthetized condition, these differences were not significant in the awake condition. In conclusion, even the richer STRF stimuli could not classify units to IC subdivisions based on response properties in the awake state. Statistically significant differences emerged only in the anesthetized state, but this could also reflect potential differential effects of anesthesia on CNIC and CtxIC responses.

In order to determine whether these statistically significant differences in FRA and STRF properties in the anesthetized condition translated into response differences to more complex stimulus types, we also acquired CNIC and CtxIC responses to an extensive stimulus battery that included two-tone stimuli, amplitude-modulated stimuli, and stimuli with embedded silent gaps in anesthetized mice (see ***Supplementary Material***). Responses to this stimulus battery were also not significantly different between CNIC and CtxIC neurons. Taken together, the FRA and STRF analyses demonstrate that CNIC and CtxIC units only show subtle differences in their responses to pure-tone and DRC stimuli in awake mice, with very few parameters showing statistically significant differences between the IC subunits. While statistically significant differences arose under anesthesia, these differences did not translate to responses to more complex stimuli.

### Single units can be reliably localized to CNIC or CtxIC by multiparametric classification of pure tone responses

Although the distributions of single FRA parameters were largely overlapping between CNIC and CtxIC units, we wondered if combining this information using machine learning tools would allow us to accomplish online determination of recording location during an experiment. We focused on FRA parameters because in contrast to STRF estimation, which is a more time-consuming and computationally intensive process, FRA parameters such as thresholds, bandwidths etc. can be rapidly measured online. To determine if recording location could be ascertained from pure-tone responses, we used three machine learning algorithms (LR, SVM, and RF; see Methods) to train classification models (with 10-fold cross validation) and compared these to the discriminative ability of single FRA parameters. Despite some statistically significant differences, single FRA parameters were ineffective at localizing recordings to IC subunits in both awake and anesthetized recordings (Fig. 6A-C). While all three types of machine learning models provided robust classification of recording location in the anesthetized state (Fig. 5D, 6B), only the RF model accomplished classification (at d’ > 1, a widely used threshold value for quantifying performance) in the awake state (Fig. 5C, 6A). As the RF model was the most successful method, we discuss this model in more detail below to aid biological interpretability.

**Figure 5:**
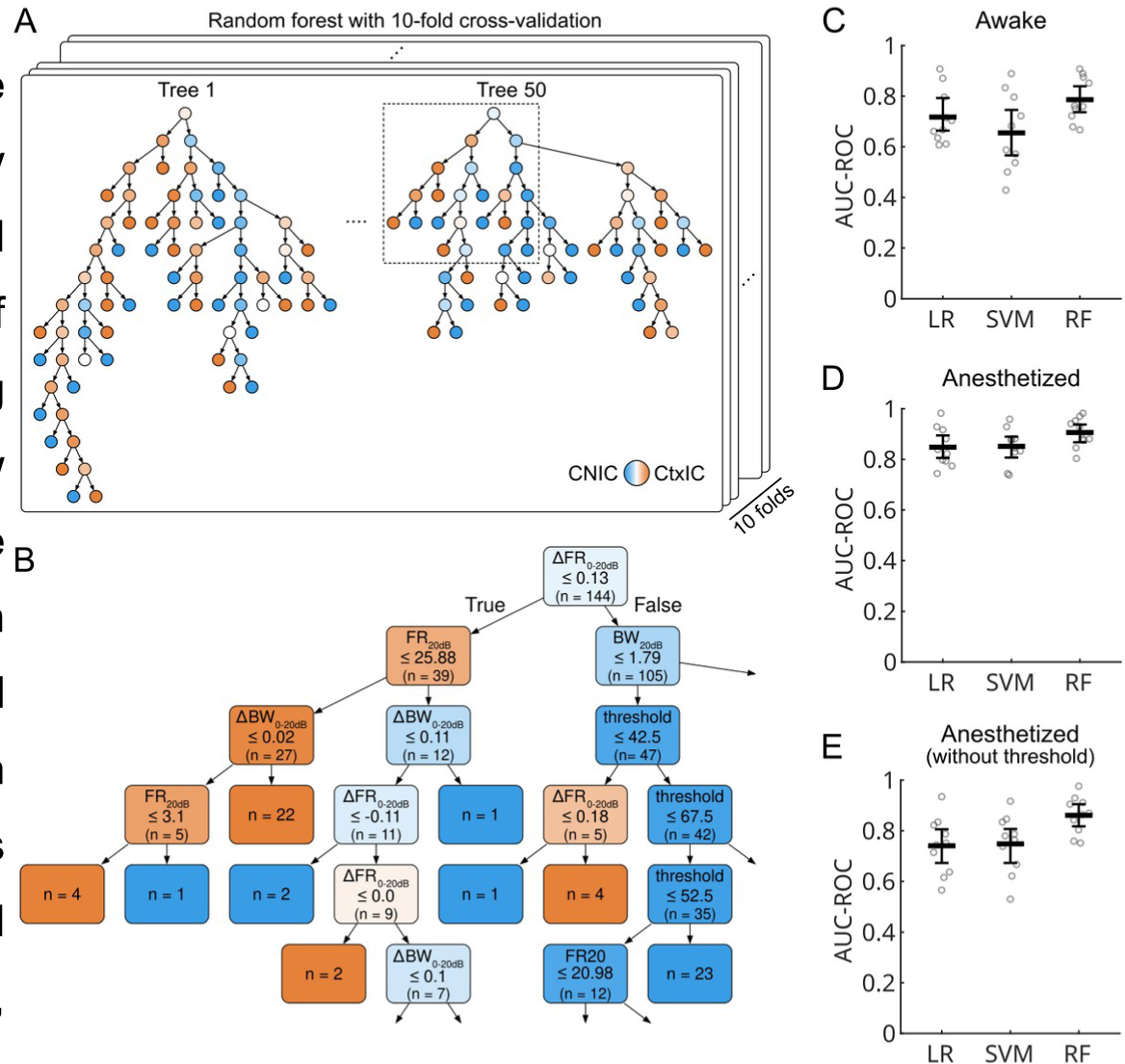
Single units can be reliably classified as CNIC or CtxIC units using parameters derived from the FRA. **(A)** Example decision trees from the RF model. Color indicates the majority class (blue – CNIC, red – CtxIC), and shading intensity indicates the strength of the majority.(**B)** An enlarged segment of decision tree 1 in A (outlined in red), showing the decision criteria for each feature, the number of input samples that meet this criteria, and the predicted output class. **(C – E)** Comparison of the performance of the three machine-learning models. Dots are area under the ROC curve plotted for each cross-validation fold. Lines are mean and standard deviation.

Broadly, RF models combine individual features of each input feature vector in a nonlinear manner using an ensemble approach to assign a category label to that input. Each of our RF ensembles was comprised of 50 decision trees (examples in Figure 5A). A decision tree model consists of internal nodes that represent predictor variables from the dataset and terminal leaf nodes that represent a distribution over the target variable, in our case, CNIC or CtxIC. A predictor variable with an assigned value is termed a feature. Each path in the tree represents a conjunction of features, and each sample in the dataset corresponds to a single path in the tree. A leaf node summarizes how many samples in the dataset correspond to that specific path as well as the majority target class of samples belonging to that path. In this way, the data set is partitioned into distinct paths. In Figure 5A, the color of the internal nodes indicates the majority class and the shading indicates the strength of the majority (e.g. dark blue indicates that most of the samples are predicted to CNIC with few CtxIC samples). The color of the terminal nodes indicates the final predicted class of the samples. By examining an example enlarged segment of a decision tree (Figure 5B), we can gain insight into which predictor variables are associated with each internal node and the target counts in the leaf nodes. The terminal leaf nodes show the number of samples that correspond to that path and list the majority target value (CNIC or CtxIC) for those samples. A predicted probability of the target class can be calculated using the proportion of test samples in the leaf node that correspond to the majority class. The RF classifier averages the predicted probabilities of the target class from all trees in the ensemble to produce an ensemble classification of the test samples.

To facilitate comparison between the machine-learning models and individual parameters, we converted the area under the ROC curve values to a sensitivity index (d’; see Methods). A d’ value greater than one is taken to indicate good model performance (typically corresponding to 76% correct). In awake animals, only the RF model achieved this threshold (Fig. 6A). In anesthetized animals, all three machine-learning models achieved d’ > 1. The FRA threshold by itself was informative at d’ > 1 in anesthetized recordings (Fig. 6B), but this likely was a consequence of the differential effect of isoflurane on CNIC and CtxIC responses. To determine the impact of this anesthetic-induced change, we re-trained our models to classify the anesthetized data while excluding the threshold as a feature. In this case, we found that similar to awake recordings, only the RF model showed performance at d’ > 1 (Fig. 6C). To determine the relative contributions of individual features to the overall performance of the RF model, we also computed the feature importances (Breiman, 2001) of the constituent individual features. In both awake and anesthetized (minus threshold) states, we found that no single feature was dominant (Fig. 6D-F). Rather, individual FRA features contributed uniformly to accomplish classification. This uniform feature importance suggests that subdivision identity is encoded in weak but distributed tuning biases rather than any single dominant response feature. As a consequence, discriminating subdivisions based on response properties was only possible by combining FRA parameters using multi-parametric, non-linear machine learning methods.

**Figure 6:**
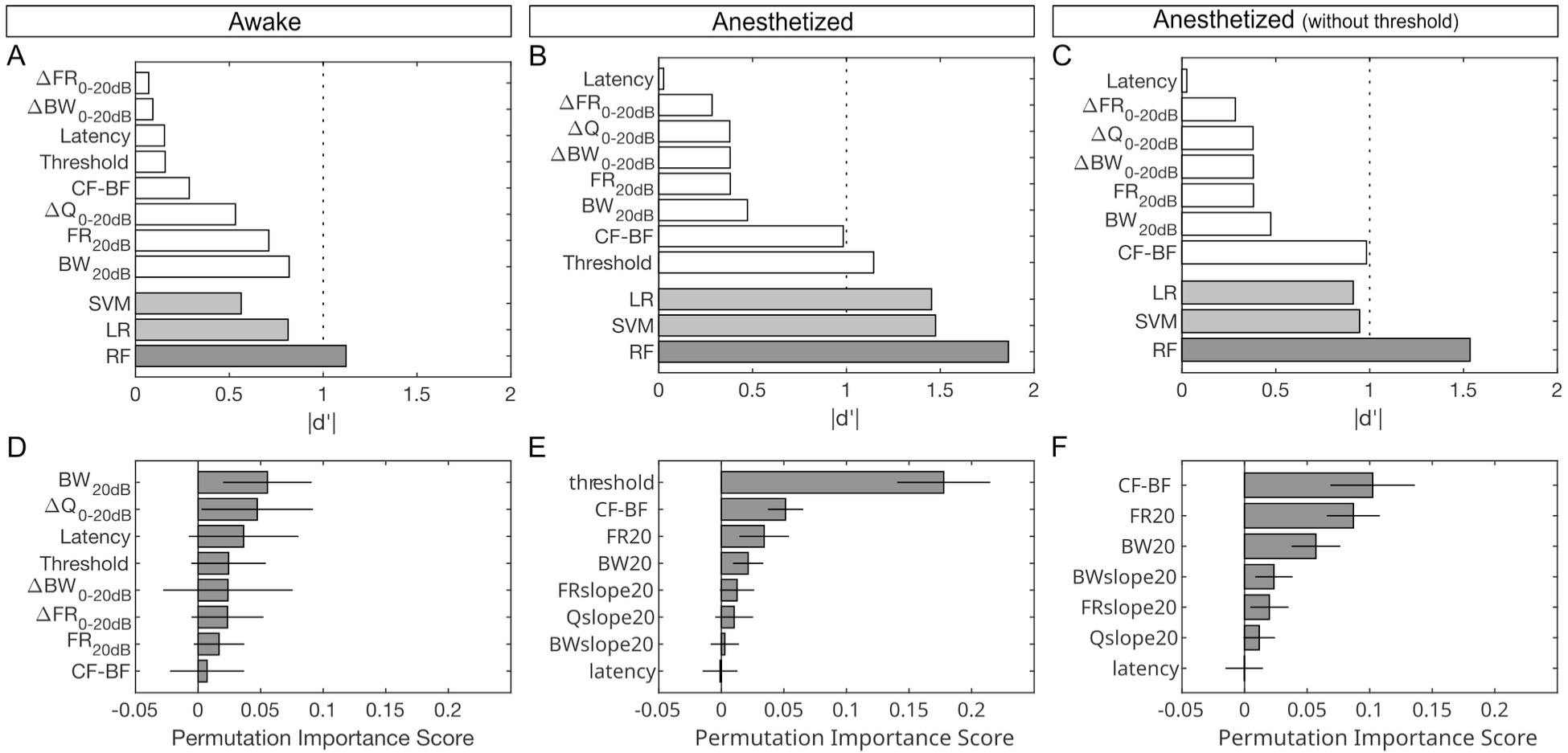
Classification performance of single parameters and machine-learning models. **(A–C)** The performance of single FRA parameters (white bars) as well as the SVM, LR, and RF models (gray bars) in correctly classifying recording location to CNIC or CtxIC in the awake and anesthetized states, and after excluding threshold as a parameter from the anesthetized state classifiers. Dashed line corresponds to a threshold of d’ = 1. **(D – F)** Feature importance scores of the FRA features that contributed to the RF model classification performance across states. Mean ± 95% CI plotted.

## DISCUSSION

The primary goal of this study was to determine whether recording location within IC subdivisions could be determined online during experiments. Online determination of location can be critical in several experimental settings, such as the injection of transplanted inhibitory precursor cells (Owoc et al., 2022), viral vectors or pharmacological agents, lesioning defined areas of interest, or microstimulation of functionally specific regions. Localization based on response properties alone would especially benefit targeting IC subdivisions in species where the IC is not close to the surface and is difficult to assess stereotactically (for example, in non-human primates) (Rocchi and Ramachandran, 2018; Slee and Young, 2011; Wang et al., 2022). Our results serve as proof-of-principle that such localization can be accomplished robustly and efficiently using responses to simple stimuli such as pure tones. Many of the above experiments are usually conducted in anesthetized animals, with isoflurane being a widely-used agent to achieve a surgical plane of anesthesia. Our results demonstrate that the machine-learning approach to achieving localization remains robust to anesthesia, further validating the applicability of this method to a range of experimental preparations. Our results also provide a potential framework accomplish response-property-based localization to other brain regions. For example, within the auditory system, distinguishing between auditory cortical subfields, especially in non-tonotopic regions such as secondary cortical areas, could potentially be accomplished based on similar techniques. Across modalities, so long as a common effective stimulus set can be presented to all recorded neurons, similar multi-parametric methods might be able to parse out neural populations even if they do not differ in their responses along single parameter axes.

At the same time, our simplifying assumption in this initial study that the IC can be treated as comprising two broad subdivisions, the CNIC and the CtxIC, warrants careful consideration. As noted in the introduction section, finer anatomical and functional parcellations exist within both of these regions. For example, the CtxIC can be subdivided into dorsal and lateral cortices (Coleman and Clerici, 1987; Druga et al., 1997; Druga and Syka, 1984; Faye-Lund, 1985; Lesicko et al., 2016; Li and Mizuno, 1997; Sitek et al., 2022; Wise and Jones, 1977), each with distinct connectivity patterns that may give rise to divergent response properties. The rostral pole of the IC has been proposed as a distinct nucleus (Meininger et al., 1986), with unique projections to the superior colliculus (Harting and Van Lieshout, 2000). Additionally, specialized neuronal populations involved in vocalization processing in bats have been reported at the ventral extent of the CtxIC (Gordon and O’Neill, 2014). Even in the absence of discrete anatomical or projection-based distinctions, response properties may vary continuously within subdivisions. Within the CNIC, for example, spatial gradients in response latency and temporal precision have been reported (Offutt et al., 2023), such that neurons in some CNIC regions exhibit response characteristics more similar to those of the CtxIC, without a clear boundary between subunits based on temporal parameters alone. Response properties within the subdivisions may also depend on molecular identity within subdivisions (Drotos and Roberts, 2024). Finally, our method for defining ground-truth boundaries using cytochrome oxidase staining may itself be imprecise at subdivision borders. Consistent with these considerations, we did not observe distinct clusters of neurons within either broad subdivision in our data, suggesting either that we over-sampled from specific finer subregions, or that response properties across finer subregions are highly variable. While we acknowledge that some of the variance observed within our broad subdivisions likely reflects this lack of finer anatomical labeling, we remain skeptical that localization at even finer spatial scales can be accomplished using single response parameters. Additionally, the spatial precision required for many experimental manipulations, such as viral delivery or microstimulation, may not necessitate this level of granularity. Therefore, our results should be viewed as a first-pass demonstration of the utility of multi-parametric approaches for online localization, which could serve as a foundation for more refined localization strategies in future studies.

Our data revealed two types of anesthesia effects. First, it greatly reduced variability in responses. For example, spontaneous rates were two-fold lower in anesthetized recordings. Maintaining a steady anesthetic plane also resulted in a more consistent internal state of the animal and reduced trial-to-trial variability of responses. These factors likely affected our estimation of some parameters. For example, spike latency could be estimated easier because of fewer spontaneous spikes. STRF predictive power was higher likely because non-auditory contributions to responses were minimal. A number of firing rate and FRA parameters may have shown significant differences between IC subunits because the distributions of these parameters under anesthesia may have been narrower. Nevertheless, our analyses demonstrated that the magnitudes of these differences remained low such that they could not drive better classification of recording location (Fig. 6). Second, anesthesia effects may have had differential effects between IC subdivisions, arising from the different effects of anesthesia on afferent and efferent pathways. While both the CNIC and CtxIC receive inputs from a variety of sources, the CNIC is the primary recipient of ascending auditory input (Beyerl, 1978; Coleman and Clerici, 1987; Shneiderman et al., 1988) while the CtxIC receives the majority of the efferent projections from the primary auditory cortex (Druga et al., 1997; Druga and Syka, 1984; Faye-Lund, 1985). We observed a strong effect of anesthesia on the FRA threshold – while thresholds spanned a broader range of values in awake animals in both CNIC and CtxIC, thresholds were consistently more elevated under anesthesia, with CNIC showing higher thresholds than CtxIC neurons. A second strong effect of anesthesia was the lack of offset responses in anesthetized recordings. We interpret this finding as an overall increase of tonic inhibition under anesthesia, which both reduced spontaneous rate (causing a floor effect that limited our ability to observe suppression below the spontaneous rate), and that reduced the amount of phasic stimulus-driven inhibition. We also observed anesthesia effects on FRA tuning symmetry (characterized using CF-BF) under anesthesia, consistent with stronger anesthetic impacts on inhibitory mechanisms.

The subtle differences we observed in the tuning bandwidth of neurons in the CNIC and CtxIC is consistent with previous reports in the cat and guinea pig (Aitkin et al., 1975; Syka et al., 2000) and may reflect the response properties of their inputs. The CNIC is the primary target of ascending projections from the cochlear nucleus (CN), superior olivary complex (SOC), and the lateral lemniscus (LL) (Beyerl, 1978; Cant, 2005; Coleman and Clerici, 1987; Oliver, 2005; Oliver et al., 1995; Riquelme et al., 2001; Shneiderman et al., 1988), while the CtxIC is the primary target of efferent and multimodal projections (Coleman and Clerici, 1987; Faye-Lund, 1985; Li and Mizuno, 1997; Oliver, 2005; Wise and Jones, 1977). The lateral CtxIC, however, receives some ascending input from the CN and sparse projections from the LL (Brunso-Bechtold et al., 1981; Cant and Benson, 2003; Coleman and Clerici, 1987; Oliver et al., 1999, 1997). The CNIC receives input from all CN subdivisions and a variety of cell types, including fusiform cells in dorsal cochlear nucleus (DCN), round and oval cells in anteroventral cochlear nucleus (AVCN), and round, oval, fusiform, and multipolar cells in the posteroventral cochlear nucleus (PVCN) (Coleman and Clerici, 1987). Conversely, the lateral CtxIC receives more restricted inputs from fusiform cells in DCN and multipolar cells in PVCN (Coleman and Clerici, 1987). While the CtxIC receives input from a limited number of CN subdivisions and cell types, they overlap with those that project to the CNIC, making it difficult to discern if these inputs contribute to the increased bandwidth we observed in CtxIC neurons compared to CNIC neurons. However, corticofugal projections from layer 5 the auditory cortex primarily target the CtxIC, and exhibit broad tuning (Williamson and Polley, 2019). Given the relative sparsity of ascending projections to the CtxIC, the tuning bandwidth of neurons in the CtxIC may be influenced primarily by efferent and intrinsic projections.

Unlike a previous study in guinea pigs (Syka et al., 2000), we did not observe latency differences between CNIC and CtxIC responses. Our estimates of spike latency (∼10 ms for both IC subdivisions in awake animals) were slightly shorter than those reported by the Syka et al. (2000) study (∼10-15 ms for CNIC and ∼15-25 ms for CtxIC), and contrary to that study, were not significantly different between IC subdivisions. One source of this difference could be methodological – Syka et al. computed the time to first spike for tones 10 dB above threshold, whereas we computed the time to significant firing rate change in the PSTH using only the highest response rates (typically at loud intensities). As onset latency varies as a function of stimulus intensity, it is expected that our study, which uses responses at louder intensities to compute latency, would show shorter latencies. A second difference could arise from differences in anesthetic use. The Syka study used urethane and ketamine as anesthetics, which could act differently on the different feed-forward regions or cell types that project to CNIC or CtxIC. The primary source of auditory inputs to the CtxIC is the ipsilateral CNIC and the auditory cortex (Morest and Oliver, 1984; Oliver, 2005), which suggests longer onset latencies in the CtxIC. But it is also known that the CN projects directly to both the CNIC and CtxIC, although whether distinct cell types (Cant and Benson, 2003) or subregions (Goldberg and Brownell, 1973) of the CN differentially project to distinct IC subregions has not been investigated in detail. Our anesthetized recordings using isoflurane yielded much shorter latencies (∼5 ms; not different between IC subdivisions), but this likely arose from better estimation of latency due to suppressed spontaneous spiking.

Regardless of anesthetic state, we found that we could classify the location of CNIC and CtxIC neurons with high accuracy based on a non-linear combination of FRA parameters. Of the tested classifiers, the RF had the best average performance, which could reflect the features of the model itself or the structure and dimensionality of the training data. LR and linear SVM algorithms both construct a single classification model with a linear decision boundary from a training dataset, whereas the RF algorithm constructs an ensemble of decision tree models using random subsets of variables from a single training dataset (Breiman, 2001). The RF algorithm uses the aggregate outputs from the collection of trees (which are nonlinear classifiers) to classify a sample, and this reduces variance without increasing overall bias, resulting in lower error and improvements in predictive performance over single models (Hastie et al., 2009). The superior performance of the RF algorithm in differentiating CNIC and CtxIC neurons may therefore be a result of the use of decision trees (which can better handle nonlinearities in data) and/or an ensemble approach. But while ensemble approaches typically increase performance, they usually do so at the expense of interpretability regarding variable interactions compared to single decision trees. However, one of the key benefits of the RF classifier is that it can rank variables by importance (as demonstrated in Fig. 6). This approach provides insight into which response properties are the most salient and how neural characteristics interact to create the diverse range of responses seen in the IC. Finally, while we have used exclusively auditory response properties to build our classifier models, it is possible that better classification can be accomplished by incorporating non-auditory information as well. Previous studies have shown that arousal state and locomotion influence IC neural activity (Yang et al., 2020; Saderi et al., 2021). Another recent study in the CtxIC demonstrated that the population response in this region is predictive of behavioral outcomes (Quass et al., 2024). Interestingly, the responses to hits and false alarms were divergent, suggesting that, on a population level, the CtxIC integrates acoustic information, motor or pre-motor activity, and behavioral outcomes (Quass et al., 2024). Although these studies did not characterize whether non-auditory influences differed across IC subdivisions, including such information in a classification scheme might further improve localization.

To our knowledge, this study is the first to classify neurons as belonging to the CNIC or CtxIC based on their response properties. Our results suggest that the location of recorded neurons can be reliably identified using relatively simple stimuli and demonstrate that appropriate machine learning techniques may be useful in characterizing neuron types in the IC. This technique could be applicable at both finer and broader levels in the future. For example, in addition to their use in discriminating neurons from the CNIC and CtxIC, another application for machine learning models could be to distinguish between neural classes within a subdivision using response properties. More broadly, our results could serve as a roadmap for classifying electrophysiological recordings from more complex brain regions that are difficult to access and distinguish using stereotactic coordinates.

## METHODS

Experimental procedures were performed in accordance with National Institutes of Health guidelines and were approved by the Institutional Animal Care and Use Committee at the University of Pittsburgh. All experiments were performed in CBA/CaJ mice (Jax 000654).

### Surgical procedures

Adult CBA/CaJ mice (6 – 13 weeks old) of either sex were sedated with an intraperitoneal injection of dexmedetomidine (IP, 0.5mg/kg) to facilitate maintenance of the anesthetic state at low doses of general anesthetic. Ten minutes after dexmedetomidine injection, general anesthesia was induced with vaporized isoflurane (2.5-3% for surgery). When subjects no longer exhibited a toe pinch reflex, they were transferred to the stereotaxic apparatus and the isoflurane lowered to a maintenance dose (2-2.5%). Body temperature was monitored and maintained at 36.5-38.5°C (FHC DC temperature controller). The IC was located using stereotaxic coordinates (-5.3 mm caudal and ± 1.3 mm lateral from bregma). Burr holes, about 1.4 mm in diameter, were drilled into the skull using a high speed stereotax-mounted drill (Model 1474, Kopf). The craniotomies were covered with 1% agarose. A custom head post was then affixed to the skull using dental acrylic (Metabond). For anesthetized recordings, the animal was transferred to the recording chamber and isoflurane anesthesia was continued for the duration of the experiment (see below). For awake recordings, we provided analgesics (Ketoprofen, 2 mg/kg) and ensure the animal was fully mobile before commencing recordings.

### Acoustic stimuli

All stimuli were generated in Matlab (Mathworks) at a sampling rate of 100 kHz, converted to analog (National Instruments), attenuated (TDT), power amplified (TDT) and delivered through a speaker (MF1, Multifield Magnetic Speaker, TDT) located approximately 10 cm from the subject on the contralateral side. Stimuli included pure tones (4 - 32 kHz), amplitude-modulated tones, two-tone complexes, tones with embedded silent gaps, and a 1-minute long segment of dynamic random chords spanning the 4 - 32 kHz frequency range (10 repetitions) to estimate spectrotemporal receptive fields (STRFs).

### Electrophysiology

All recordings were conducted in a double-walled sound-attenuating booth (IAC), the walls of which were lined with anechoic foam (Pinta Acoustics). Subjects were headfixed to a vibration-isolation tabletop. For anesthetized recordings, isoflurane anesthesia (0.5-1.5%) was provided through a face mask and body temperature maintained at 36.5 – 38.5°C (FHC DC temperature controller). For awake recordings, animals were placed in a custom restraint, and a piezoelectric sensor used to monitor motion. Single units were recorded using a 64-channel silicon probe (64D sharp, fabricated by IDAX Microelectronics) (Du et al., 2011) with individual sites electroplated with nanogold particles to achieve an impedance of ∼2 – 3 MΩ. The electrode was dipped in DiI to allow for post-hoc verification of the recording site. The agarose was removed from the craniotomies and the probe positioned above the IC at a 15° angle, pointed rostrally, and lowered to a depth of 1.5 mm using a hydraulic microdrive (FHC Inc.). Mineral oil was placed over the craniotomy sites to prevent the tissue from drying. Electrophysiological signals were digitized and amplified using a low-noise amplifier (Ripple Scout) and visualized using Trellis suite software (Ripple, Inc.). The data was sorted off-line to isolate single unit clusters using JRCLUST (for anesthetized recordings; Jun et al., 2017) or Kilosort4 and Phy (for awake recordings, Pachitariu et al., 2024). Cluster quality was assessed using a range of metrics (e.g. signal-to-noise ratio of spike waveform, ISI histogram, stability of cluster over recording duration). Only well-isolated single unit clusters (SNR > 4) with low refractory period violations (<10%) and which had consistent firing rates throughout the presentation of a stimulus set were considered for analysis. Post-hoc reconstruction of the probe track was used to obtain the ground-truth location of recorded single units in the IC. Multiunit activity (MUA) was used to determine range of frequencies represented along the probe track. MUA was defined as the envelope of the band-pass filtered (350-3500Hz) and rectified raw signal, in response to pure tones of different frequencies (6, 8, 16, and 24kHz).

### Post-hoc verification of probe location

At the end of the recording session, we performed histochemistry for cytochrome oxidase to delineate the borders of the CNIC anatomically (Fig. 1B,C) (Cant and Benson, 2005; Ito et al., 2018) and used the DiI or DiO staining to register the probe trajectory to the underlying IC anatomy. Subjects were transcardially perfused with 0.1M phosphate buffer saline (PBS) followed by 4% paraformaldehyde (PFA) in 0.1M PBS. The brains were extracted and post fixed in 4% PFA overnight at 4°C. Following post fixation, the brains were cryoprotected in 30% sucrose in 0.1M PBS for 48 hours at 4°C. Coronal sections (50µm in thickness) were cut using a freezing microtome. Sections were washed in 0.1 M PBS then incubated at room temperature in a solution containing 10 mg diaminobenzidine (DAB, Sigma D5905), 20 mL 0.1M PBS, 5mg cytochrome oxidase (Sigma C2037), and 0.8 g sucrose. Sections were incubated at room temperature until the CNIC appeared well differentiated (typically 4 hours). Sections were washed three times in 0.1 M PBS and mounted onto superfrost plus microscope slides (Fisherbrand). Light and fluorescent microscopy was performed to visualize the DAB reaction product (Figure 1A) and DiI or DiO, respectively. Digitized images of coronal sections were obtained (Axio Imager, Zeiss) and overlaid in ImageJ. The boundary of the CNIC was then determined. Probe depth was adjusted for post-fixation brain tissue shrinkage by a factor of 0.88, a value established in separate experiments independent of the primary recording sessions. The depth of each unit was defined as the location of the maximum peak-to-trough amplitude of mean waveform templates. Further subdivision of the CtxIC into the dorsal and lateral cortices was not performed.

### Data analysis – tone response parameters

The frequency response area (FRA) was characterized by acquiring responses to 100 ms pure tones (Inter-stimulus interval of 300 ms) ranging from 4 to 32 kHz (10 steps/oct.) at various sound levels (25 – 80 dB SPL for awake mice and 35 – 80 dB SPL for anesthetized mice, 5 dB spacing) presented in pseudo-random order. The spontaneous firing rate was determined during the 100 ms segment of the inter-stimulus interval immediately preceding stimulus onset. The onset/offset ratio was determined by as the firing rate in a 0 – 50 ms window after sound onset divided by the firing rate in a 0 – 50 ms window after sound offset. The best frequency (BF) was defined as the frequency that elicited the highest firing rate at any sound level. Threshold was defined as the lowest sound level to evoke a response significantly different from baseline (i.e., greater than mean + 2 SD of spontaneous rate). The characteristic frequency (CF) was defined as the frequency that elicited the highest firing rate at threshold. Onset latency was determined using the peristimulus time histogram (PSTH, 1 ms bins) of the most active trials (≥90^th^ %ile). A piecewise linear fit was applied to the PSTH (from -100 ms to +50 ms relative to sound onset) using maximum likelihood estimation (`mle`, MATLAB) with a breakpoint constrained between 3 ms and 50 ms. The estimated breakpoint was defined as the latency. Bandwidths were defined as the frequency ranges where driven firing rates significantly exceeded baseline. For discontinuous frequency tuning curves, the bandwidths of individual segments were summed. For each neuron, linear fits of the bandwidth in octaves and firing rates between the threshold and 20 dB above threshold were used to estimate the slopes of the bandwidths and firing rates in this sound level range. The quality factor Q, defined as the best frequency divided by the bandwidth in Hz, was calculated at each level relative to threshold.

### Data analysis – spectrotemporal receptive fields

Spectrotemporal receptive fields (STRFs) were estimated using NEMS (Pennington and David, 2020; Thorson et al., 2015) and previously described methods (Montes-Lourido et al., 2021). Briefly, the actual peristimulus time histogram (PSTH) of unit responses to dynamic random chords was computed in 5 ms bins, averaged over 10 repetitions. Nested cross-validation, where 90% of the data was used to fit the model and the remaining 10% was used to test the model, was repeated 10 times using non-overlapping segments to yield 10 STRF estimates which were then averaged to result in a mean STRF. A significance mask was calculated using a permutation test (Montes-Lourido et al., 2021), and non-significant STRF weights were set to zero. The correlation coefficient between the predicted responses to the test data sets and actual responses (r-test) was used as a metric of goodness of fit. STRFs with an r-test <0.3 were excluded from further analysis. The temporal and frequency response profiles of the average STRF estimate for each neuron were estimated using a Gaussian fit (Matlab, MathWorks).

### Classifier models

Logistic regression (LR), linear support vector machine (SVM), and random forest classifiers (RF) were trained on pure-tone responses (used to derive the FRA) to determine whether CNIC and CtxIC responses could be reliably separated based on FRA parameters. Models were trained and evaluated using scikit-learn (Pedregosa et al., 2011). Missing values were estimated using SimpleImputer (Scikit-learn toolbox in Python). Anatomical probe location ground-truth, determined using DiI or DiO and cytochrome oxidase labeling, was used to create a binary target variable. Further details of data used for classification are shown in Table 1. We used 10-fold nested cross validation, where 90% of non-overlapping data segments were used to fit the model and the remaining 10% was used to test the model with each segment containing an equal proportion of CNIC and CtxIC units, repeated 10 times. The LR algorithm was implemented with the “liblinear” solver. The SVM algorithm used a linear kernel. The RF algorithm was implemented with an “entropy” scoring criterion and an ensemble size of 50. Model performance was evaluated based on the area under the receiver operating characteristic (ROC) curve, a measure of discrimination. Median accuracy and area under the ROC curve was calculated across the 10 runs for each type of model. Model discriminability was calculated by approximating d’ from the area under the ROC curve (AUC) using the equation 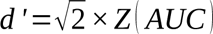, where *Z* (*x*) is the inverse of the cumulative normal distribution. Model discriminability was then compared to the discriminability achieved using each individual training variable (Table 1) by calculating the absolute value of d prime (|d’|), the standardized difference between the means of the CNIC and CtxIC distributions 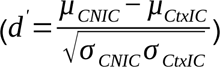.

### Feature importances

The importance of features used for classifier models was evaluated using the permutation method (Breiman 2001; using `permutation_importance,` Scikit-learn toolbox in Python). Permutation importance was defined as the difference between AUC from all features and AUC from permuting a feature. We generated 10 scores for each feature (corresponding to 10 cross-validation folds per classifier; means shown in Fig. 6).

### Statistics

Statistical analysis was performed using MATLAB. Data were evaluated for normal distribution using a one sample Shapiro-Wilk test. Two sample t-tests or Mann Whitney tests were used to compare two independent groups with either normally or non-normally distributed data, respectively. One-way ANOVA or Kruskal-Wallis tests were used to compare three or more groups with either normally or non-normally distributed data, respectively. Multiple comparisons were corrected for using the Dunn pairwise method. Statistical significance for all tests was set to p<0.01 and corrected for in cases of multiple comparisons.

## DATA AVAILABILITY

The data that support the findings of this study are available from the corresponding author upon reasonable request.

## CODE AVAILABILITY

Custom analysis code used in this study is available from the Sadagopan lab’s Github repository: https://github.com/vatsunlab/CNIC_CtxIC

## ACKNOWLEDGEMENTS

This work was supported by the National Institutes of Health [F30DC018185, R01DC004199, and R01DC019814] and the CURE Phase 17 Formula Funds from Commonwealth of PA Dept. of Health. The funding sources had no role in study design, data collection or analysis, publication, or preparation of the manuscript.

## AUTHOR CONTRIBUTIONS

**MSO:** Conceptualization, anesthetized experiments, formal analysis, writing. **JL:** Awake experiments, data curation, formal analysis. **AJ:** Formal analysis. **KK:** Conceptualization, supervision, funding acquisition. **SS:** Conceptualization, software, writing, supervision.

## COMPETING INTERESTS

The authors declare no competing interests.

## Notes

### Competing Interest Statement

The authors have declared no competing interest.

