## Supplementary Information for "Multiparametric Classification of Pure-tone Responses Distinguishes Neurons in Inferior Colliculus Subdivisions"

### SUPPLEMENTARY RESULTS

#### *STRF observations suggest putative differences may emerge between CNIC and CtxIC responses for complex stimulus types*

In both CNIC and CtxIC, E frequency tuning widths determined from STRFs (awake CNIC median = 0.11 oct., CtxIC median= 0.17 oct.) were substantially and significantly ($p < 0.0001$) narrower than bandwidths computed from the FRA, even at 20 dB above threshold (awake CNIC median= 0.60 oct., CtxIC median= 0.8 oct.). This difference persisted across both the awake and anesthetized states. These observations suggested that the denser DRC stimulus might have engaged side-band suppression, which was not activated by single tones. The very low spontaneous rates in our anesthetized preparation may have additionally hampered our ability to observe suppression in single-tone (FRA) responses. This led us to wonder whether additional differences might emerge between CNIC and CtxIC units in terms of different levels of sideband suppression. Similarly, our STRFs revealed complexities in the temporal response profiles of neurons in both the CNIC and CtxIC in both the anesthetized and awake states, which led us to wonder if additional differences might emerge if temporally varying stimuli were employed. To investigate these possibilities, we presented a stimulus battery consisting of two-tone stimuli, amplitude modulated stimuli, and stimuli with embedded silent gaps. Due to the extended recording stability required for completing this exhaustive stimulus set, we acquired these data only in the anesthetized state.

#### *CNIC and CtxIC neurons exhibit similar amounts of two-tone suppression in anesthetized mice*

We first examined the effects of two simultaneously presented pure tones on CNIC and CtxIC response rates in anesthetized mice. By positioning one tone at the estimated best frequency, we reasoned that we could drive units robustly so that suppression by a second frequency could be observed. Because of our array recording set up, the exact best frequencies of units could not be estimated until units were sorted offline after the experiment. Therefore, during the experiment, we presented two tones by fixing one of the tone frequencies (F1) at a few values that spanned the range of frequencies observed in a given track. The second tone frequency (F2) was roved in a ± 0.5 to 2 octave range relative to F1. For analysis, the best frequency of each unit was approximated to be the frequency that elicited the highest response when the two frequencies were equal (F1=F2). If the strongest response rate was not at F1=F2, it was likely that we did not accurately position F1 at the best frequency during the experiment, and these data were not used.

A subset of units (CNIC n=16/108 and CtxIC n=11/107) demonstrated a response that was greatest at F1=F2. In both CNIC and CtxIC units, we saw a variety of responses, including units with sustained side-band suppression (Fig. S1A, left) and those with variable suppression (Fig. S1A, right). On average, units demonstrated a maximum firing rate suppression of 67% ± 28.1% in the CNIC and 75% ± 10.7% in the CtxIC. There was no significant difference in the maximal percent reduction between the two populations (p= 0.39, unpaired t test; Fig. S1B). Next, we quantified total suppression as the area under the curve from -0.5 to 0.5 oct and -1 to 1 oct (Fig. S1C). This analysis can differentiate between sustained suppression and variable or narrow suppression, even if two units exhibited the same maximal suppression. We did not observe significant differences in the area under the curve between -0.5 and 0.5 oct (CNIC mean= 0.46, SD= 0.25; CtxIC mean= 0.37, SD= 0.12; p= 0.27, unpaired t test) or -1 and 1 oct (CNIC median= 0.95, IQR= 0.75; CtxIC median= 0.90, IQR= 0.4; p= 0.72, Mann-Whitney test, Fig. S1C). Together, these data suggest that while both CNIC and CtxIC units are indeed influenced by side-band suppression, they exhibit similar degrees of suppression and this parameter cannot be used to discriminate recording location. We note again that these analyses are constrained by our array recording methodology, in that we could not present stimuli with one tone precisely at the BFs of recorded units. Additionally, because we approximated BF as the frequency at which the maximal response was observed at F1 = F2, we could not observe two-tone facilitation.

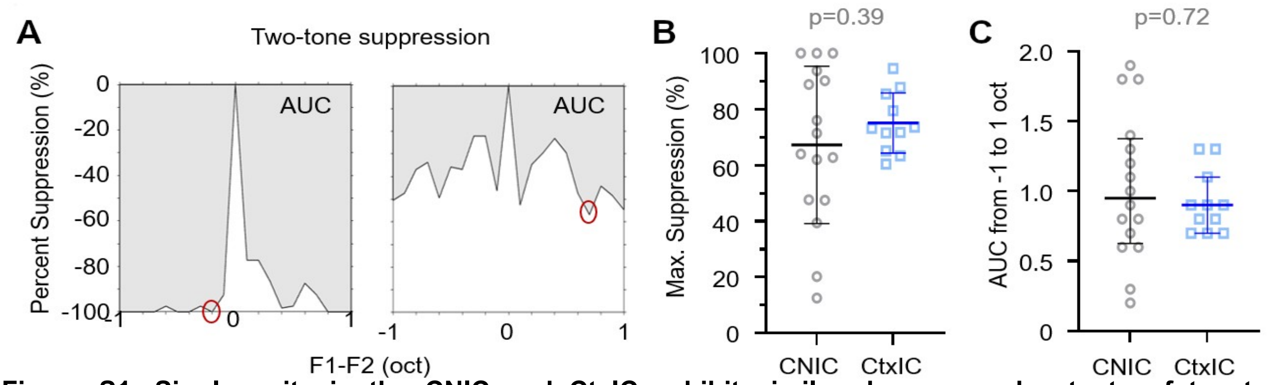


**Figure S1. Single units in the CNIC and CtxIC exhibit similar degrees and extents of two-tone suppression. (A)** Representative two-tone suppression responses from two units from the CNIC. Best frequency was approximated as the frequency at which the peak response occurred when F1=F2. The left example illustrates strong sideband suppression (shaded region) over a large range of frequencies. The right example shows more variable suppression with multiple peaks and valleys. From the two-tone suppression plots, we extracted the maximal percent suppression (red circles), and area under the curve (AUC, shaded regions). **(B)** The degree of maximal percent suppression was similar between CNIC and CtxIC units. There was no significant difference between the populations (p= 0.39, Unpaired t test). **(C)** Total sideband suppression (AUC) from -1 to 1 oct was similar between CNIC and CtxIC units (p= 0.72, Mann-Whitney test).

***The majority of IC single units are gap detecting, with similar gap detection thresholds in CNIC and CtxIC units***

We next presented a series of stimuli to further probe the temporal integration properties of CNIC and CtxIC units. First, we determined neural response types and gap detection thresholds by presenting pure tones with embedded gaps. Example post-stimulus time histograms (PSTHs) of CNUC responses to these stimuli are shown in Fig. S2A. We found that the majority of units (CNIC= 66%, CtxIC= 73%) detected gaps at one or more tested frequencies (Fig. S2B, top). Next, we classified single units according to three main response types: onset, sustained, and offset. Units that did not fit into these three categories were classified as ‘Other’. As with the two-tone data, due to our array recording approach, the response type was determined based on an approximation of the best frequency, which we estimated as the frequency that elicited the highest firing rate in the absence of a gap. We acknowledge that this strategy might bias our analyses in favor of onset and sustained responders. In addition to characterizing the response type using the objective measures described in the methods section, each response was also manually inspected and only one unit displayed an offset response and this response type was observed at a frequency other than the estimated best frequency. Thus, our single units were classified as onset, sustained, or other (Fig. S2A). Single units classified as “other” responders typically had inconsistent response types that varied between trials, typically between onset and sustained response types. The percentage of single units in the onset, sustained, and other categories were roughly equal in both the CNIC and CtxIC (Fig. S2B, bottom).

We then investigated the gap detection threshold (GDT) at the estimated best frequency (est. BF) of single units in the CNIC (Fig. S2C, left) and CtxIC (Fig. S2C, right) according to response type. We found that the GDT at est. BF appeared to vary with response type. The lowest GDTs were observed in single units with a sustained response type. Additionally, sustained responders were the least likely to have no GD. Conversely, roughly 40% of onset responders, in both the CNIC and CtxIC, exhibited no GD. Onset responders that did exhibit GDTs responded to gap length greater than or equal to 5ms. Single units that were classified as “other” responders, which displayed mixed response types, were the most likely to have no GD, regardless of recording location. While there appears to be a relationship between response type and GDT, the relatively equal distributions of response types in our single units in the CNIC and CtxIC (Fig. S2B, bottom) allowed us to compare the GDT at est. BF across all units, regardless of their response type. We found that the GDTs at est. BF were distributed across all tested gap lengths in both the CNIC and CtxIC (Fig. S2D). Furthermore, there was no difference in the cumulative distribution of GDTs between the two groups ($p= 0.64$, Kolmogorov-Smirnov test). Thus, single units in the CNIC and CtxIC exhibited similar distributions of response types and GDTs at the estimated BF.

Next, to investigate the relationship between the GDT and carrier frequency, we examined the GDT at each carrier frequency relative to the est. BF (Fig. S2E). We found that, in both the CNIC and CtxIC, the GDT was lowest for carrier frequencies at or near the est. BF, and increased with increasing distance from the est. BF. To confirm this, we averaged the GDT of single units at frequencies ±0.8 oct relative to the est. BF and fit quadratic functions to the average GDT of single units in the CNIC and CtxIC. This frequency range was chosen because the majority of our single units displayed GDT in this range. The quadratic fit of the average GDTs in the CNIC and CtxIC confirmed that the minimum GDT was observed at or near the est. BF with increasing GDTs in the sidebands (Fig. S2E). However, while 57% (48/84 single units) of CNIC units exhibited a minimum GDT at est. BF, 43% (36/84 single units) had a minimum GDT in the sidebands. In the CtxIC, 38% (37/98 single units) of single units exhibited a minim GDT at est. BF while 62% (61/98 single units) had minimum GDTs at frequencies greater than the est. BF. These data suggest that while on average the GDT is lowest at or near the est. BF in both the CNIC and CtxIC, there may be differences in how individual units from these populations respond to offset carrier frequencies.

To further investigate gap detection in offset carrier frequencies, we determined 1) the difference between the est. BF and the minimum gap detection frequency and 2) the difference between the GDT at est. BF and the minimum GDT. When comparing the cumulative distributions (Fig. S2F, inset) of the difference between the minimum gap detection frequency and the est. BF of single units in the CNIC and CtxIC, we found a significant difference ($p= 0.009$, Kolmogorov-Smirnov test), with single units in the CtxIC more likely to exhibit a minim GDT in the sideband than single units in the CNIC. Additionally, we found that the cumulative distributions (Fig. S2G, inset) of the difference between the GDT at est. BF and minimum GDT of single units in the CNIC and CtxIC were significantly different ($p= 0.046$, Kolmogorov-Smirnov test), with single units in the CtxIC more likely to exhibit a larger difference in the GDT at est. BF and minim GDT than single units in the CNIC. Together, these data suggest that units in the CtxIC are more likely to exhibit improved gap detection in offset carrier frequencies than units in the CNIC.

In summary, our data indicate that single units in the CNIC and CtxIC have similar gap detection capabilities, with the majority of units responding to silent gaps in pure tones and similar distributions in their response types and gap detection thresholds at the est. BF. Interestingly, we found that units in the CtxIC were more likely to have minimum GDT in the sideband than units in the CNIC. Additionally, units in the CtxIC were more likely to have a larger difference in their minimum GDT and GDT at est. BF. While the frequencies tested here are too far apart to truly characterize gap detection in the sideband, these data suggest that a subset of IC neurons may have improved gap detection in the sideband and warrants further investigation.

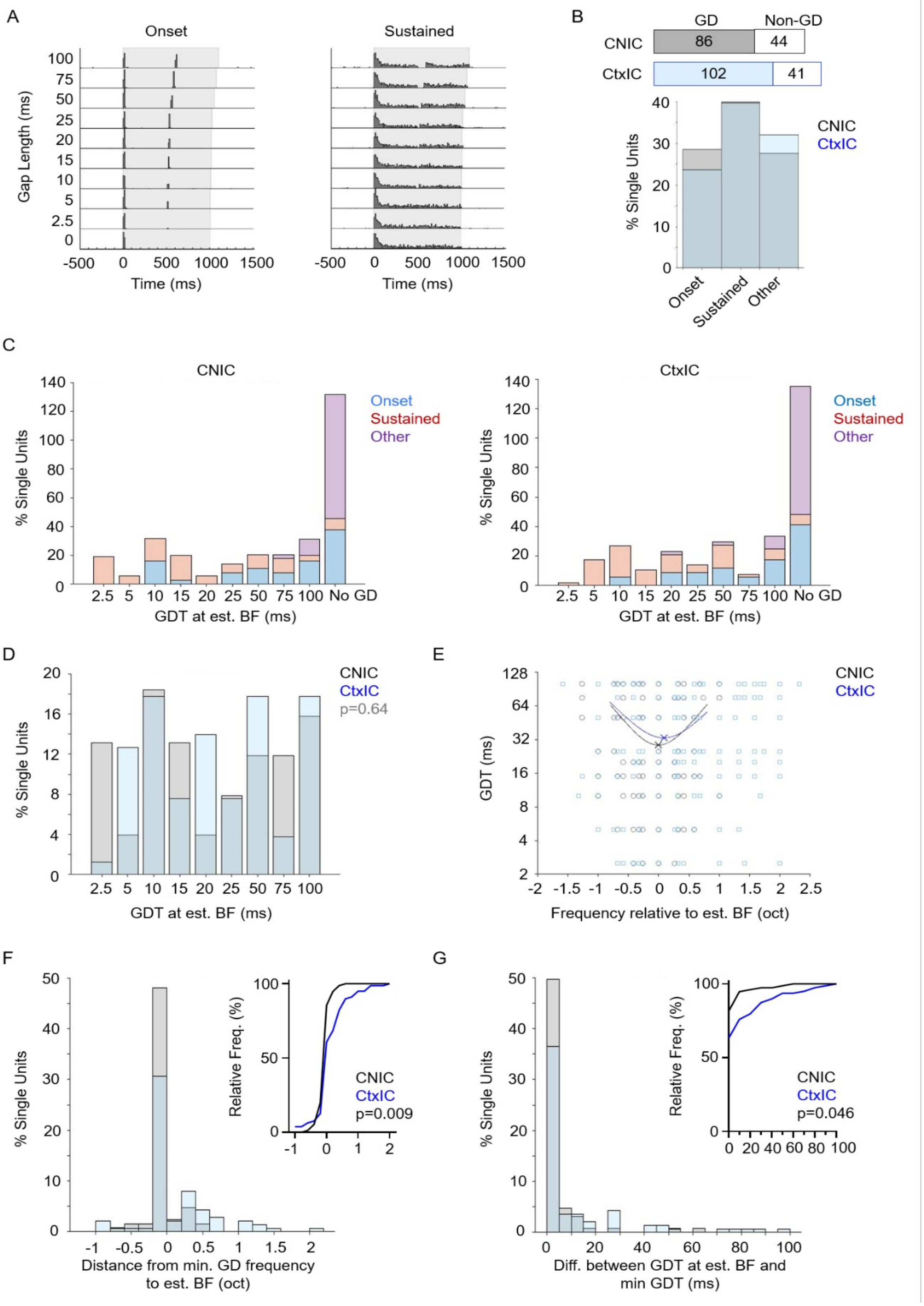
A
Onset
Sustained
Gap Length (ms)
100
75
50
25
20
15
10
5
2.5
0
-500 0 500 1000 1500
Time (ms)
B
GD
Non-GD
CNIC 86 44
CtxIC 102 41
% Single Units
CNIC
CtxIC
Onset
Sustained
Other
C
CNIC
CtxIC
Onset
Sustained
Other
% Single Units
2.5 5 10 15 20 25 50 75 100 No GD
GDT at est. BF (ms)
D
% Single Units
CNIC
CtxIC
p=0.64
GDT at est. BF (ms)
E
GDT (ms)
CNIC
CtxIC
Frequency relative to est. BF (oct)
F
% Single Units
Relative Freq. (%)
CNIC
CtxIC
p=0.009
Distance from min. GD frequency
to est. BF (oct)
G
% Single Units
Relative Freq. (%)
CNIC
CtxIC
p=0.046
Diff. between GDT at est. BF and
min GDT (ms)

**Figure S2. Gap detection thresholds and response types of CNIC and CtxIC units. (A)** Examples of single units from the CNIC showing onset (left) and sustained (right) responses. **(B)** *Top,* Relative prevalence of gap detecting (GD) units in the CNIC (grey) and CtxIC (blue). *Bottom,* The distribution of gap response types at the estimated best frequency of units in the CNIC and CtxIC. **(C)** Histograms of the gap detection threshold (GDT) at the estimated best frequency (est. BF) of single units displaying different response types (onset – blue, sustained – red, other - magenta) in the CNIC (left) and CtxIC (right). Sustained responders typically displayed the lowest GDTs. **(D)** Histograms of the GDT at the est. BF of single units in the CNIC (grey) and CtxIC (blue). There was no difference in the cumulative distributions of GDTs between the two groups (p= 0.64, Kolmogorov-Smirnov test). **(E)** GDT at each carrier frequency relative to the est. BF. The GDT was typically lowest for carrier frequencies at or near the est. BF of single units in both the CNIC (grey) and CtxIC (blue). Lines correspond to a quadratic fits, the est. BF at which the minimum GDT was observed is marked with an "X". **(F)** Histograms of the difference (in oct.) between the minimum gap detection frequency and the est. BF of single units in the CNIC (grey) and CtxIC (blue). *Inset,* cumulative histograms. CNIC and CtxIC distributions were significantly different (p= 0.009, Kolmogorov-Smirnov test), with single units in the CtxIC more likely to exhibit a minimum GDT in offset carrier frequencies than single units in the CNIC. **(G)** Histograms of the difference (in ms) between the GDT at est. BF and minimum GDT of single units in the CNIC (grey) and CtxIC (blue). *Inset,* cumulative histograms. CNIC and CtxIC distributions were significantly different (p=0.046, Kolmogorov-Smirnov test), with single units in the CtxIC more likely to exhibit a larger difference in the GDT at est. BF and minimum GDT than single units in the CNIC.

***The majority of single units in the CNIC and CtxIC responded to amplitude modulated tones, with the temporal best modulation frequency (tBMF) lower than the rate-based best modulation frequency (rBMF)***

In addition to gap detection thresholds, we assessed temporal processing by characterizing CNIC and CtxIC responses to sinusoidal amplitude modulated (sAM) tones. The majority of single units responded to sAM tones, with similar responses in the CNIC and CtxIC. We observed both sustained (Fig. S3A) and onset (Fig. S3B) responses in both IC subunits. Significant synchrony, or temporal modulation of firing rate, was observed in 73% (82/113 single units) of CNIC units and 60% (79/131 single units) of CtxIC units at one or more carrier frequencies (Fig. S3C, *top*). CNIC units had a median tBMF of 12Hz while CtxIC units had a median tBMF of 14Hz (Fig. S3C, *bottom*). There was a significant difference in the cumulative distributions of the tBMF in CNIC units compared to CtxIC (p= 0.01), with CtxIC units more likely to have a higher tBMF than CNIC units (Fig. S3C, *inset*). Significant modulation of response rate as a function of sAM frequency was observed in 89% (101/113 single units) of CNIC units and 86% (113/131 single units) of CtxIC units at one or more carrier frequency (Fig. S3D, *top*). The distribution of the rBMF was skewed towards higher AM frequencies for both CNIC and CtxIC units, with a median rBMF of 87Hz in CNIC units and 66Hz in CtxIC units (Fig. S3D, *bottom*). There was no significant difference in the cumulative distributions of the rBMF in CNIC and CtxIC (p= 0.22, Fig. S3D, *inset*). Together, these data suggest that while CNIC neurons demonstrate an increased propensity for synchronized responses, CtxIC neurons may have improved synchrony at higher temporal modulation frequencies.

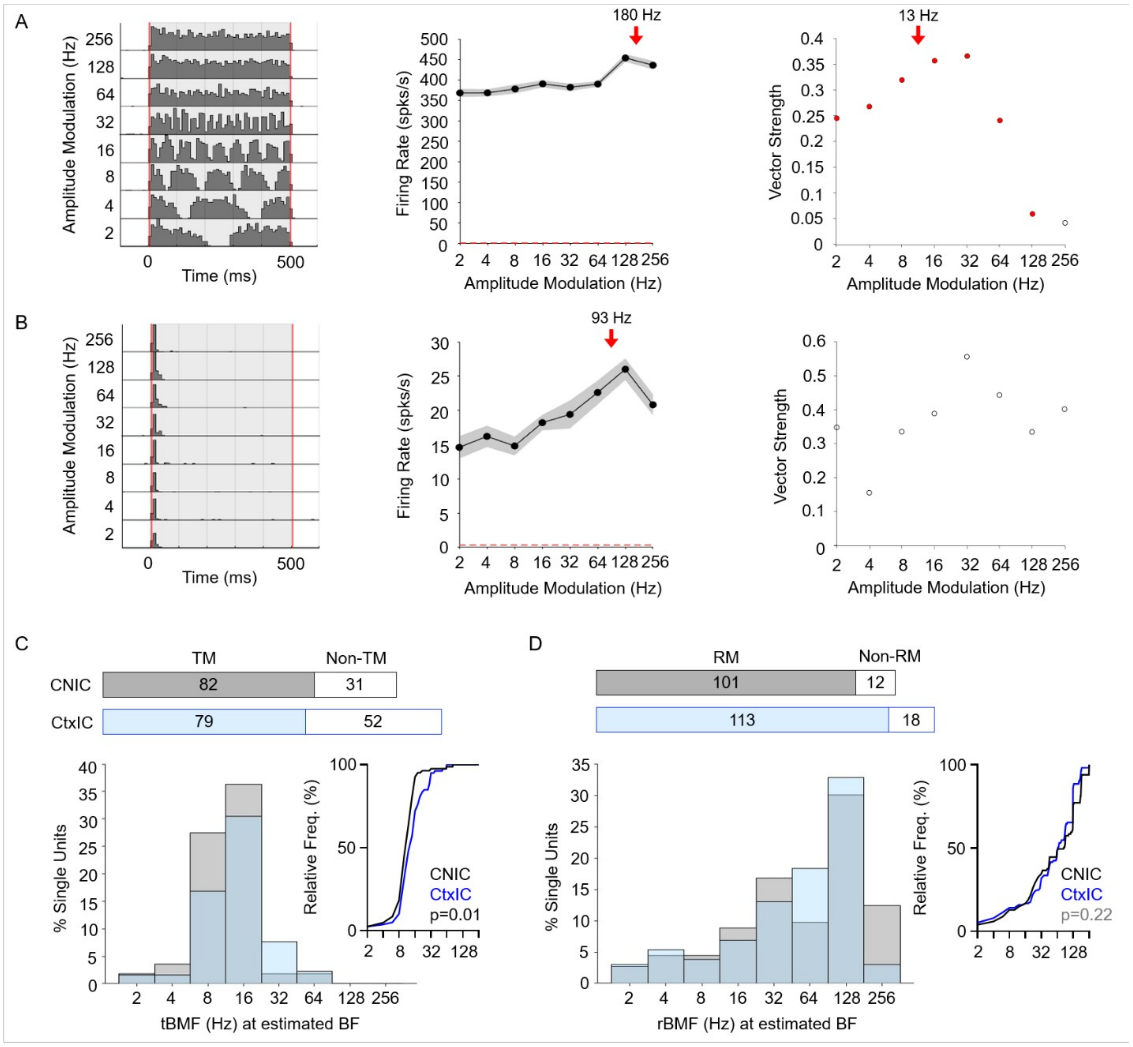


**Figure S3. Responses of CNIC and CtxIC units to sinusoidally amplitude modulated tones. (A)** Example PSTH (left) of a CNIC unit with both rate based (middle) and temporal based (right) response modulation. Rate based modulation (middle) was characterized as an increase in the firing rate at one or more AM frequency, the rBMF of this unit was 180 Hz (red arrow). Temporal based modulation (right) was exhibited at AM frequencies with a significant vector strength (Rayleigh statistic $\geq$ 13, red filled dots), the tBMF for this unit was 13 Hz (red arrow). **(B)** Example PSTH (left) of a CNIC unit with rate based (middle), but not temporal based (right), reponse modulation. The rBMF of this unit was 93 Hz (middle, red arrow). This unit did not exhibit significant temporal modulation (right) at any AM frequency (Rayleigh statistic < 13, open circles). **(C)** Top, the majority of single units temporally modulated their firing rates in response to sAM stimuli, with 73% (82/113 single units) of CNIC units and 60% (79/131 single units) of CtxIC units reaching a significant vector strength at one or more carrier frequency. Bottom, Histograms (bottom) of the tBMF at the estimated best frequency (BF) of single units in CNIC (grey) and CtxIC (blue). The cumulative distributions (inset) of tBMF at the estimated BF demonstrates that CtxIC units were more likely to exhibit a higher tBMF compared to CNIC units (p=0.01, Kolmogorov-Smirnov test). **(D)** *Top,* the majority of single units also exhibited overall firing rate changes as a function of sAM frequency, with 89% (101/113 single units) of CNIC units and 86% (113/131 single units) of CtxIC units demonstrating significant rate modulation at one or more carrier frequency. *Bottom*, the distributions of rBMF were skewed toward higher AM frequencies in both the CNIC (grey) and CtxIC (blue). There was no significant difference in the cumulative distributions (inset) of the rBMF in CNIC and CtxIC (p= 0.22, Kolmogorov-Smirnov test).

### SUPPLEMENTARY DISCUSSION

Our supplementary results should be interpreted while keeping in mind a methodological limitation. While recording using high-density multielectrode arrays enabled us to simultaneously record the activity of a large number of neurons, we could reap the benefits of this technique only when using fixed stimulus sets (such as the FRA and STRF data in the main manuscript). But when stimuli needed to be tailored to a specific neuron's preferences – for example, for choosing the carrier frequency for tone AM stimuli or for fixing one frequency at the BF for two-tone stimuli – array recordings were not optimal. In our supplementary experiments and analyses, we approximated the best frequency for two-tone, gap detection, and MTF analysis because we could not assess individual neuron BFs online. This affected our interpretation of results from the gap detection, amplitude modulation, and two-tone experiments. Indeed, this factor may have contributed to why we observed temporal receptive field differences between the STRFs of CNIC and CtxIC neurons that did not extend to gap-detection and two-tone stimulus sets (in anesthetized mice).

Although temporal tuning differences estimated from the STRF did not translate to other commonly used measures of temporal processing, such as gap detection thresholds or temporal and rate-based modulation response to AM tones, our data do show hints of differences in temporal processing in the sideband. For example, CtxIC neurons were more likely to have a minimum gap detection threshold at a non-best frequency carrier tone compared to CNIC neurons. Additionally, the difference between the gap detection threshold at the estimated best frequency and the minimum gap detection threshold was more likely to be greater in CtxIC neurons than CNIC neurons. While the frequencies tested here were too coarsely spaced to characterize gap detection in the sideband in detail, these data suggest that a subset of IC neurons, particularly in the CtxIC, may have improved gap detection at non-preferred carrier frequencies. However, as previously stated, isoflurane anesthesia suppresses auditory cortex activity, and recent research suggests that cortico-collicular projections enhance gap detection in the dorsal IC (Weible et al., 2020). As the primary recipient of cortico-colliular projections (Druga et al., 1997; Druga and Syka, 1984; Faye-Lund, 1985), the CtxIC would be most affected by cortical suppression. Thus, cortical suppression may lead to increased gap detection thresholds in CtxIC neurons at the best frequency while offset carrier frequencies remain unaffected. This could explain our finding CtxIC neurons were more likely to have minimum gap detection thresholds at offset carrier frequencies.

In response to AM tones, we observed similar temporal and rate-based modulation response in CNIC and CtxIC neurons. Interestingly, we found that a higher fraction of CNIC than CtxIC neurons demonstrated temporally modulated responses to AM stimuli; although CtxIC neurons were more likely to have a higher tBMF than CNIC neurons. Previous research suggests that both the rate and temporal-based modulation in the CNIC is stimulus dependent and varies based on the carrier frequency and sound pressure level (Krishna and Semple, 2000). However, because we observed rate and temporal-based modulation at multiple carrier frequencies (Krishna and Semple, 2000), it is rather unlikely that the higher number of CNIC neurons exhibiting temporally modulated responses to AM tones we encountered reflects an effect of the carrier frequencies. If further research confirms this finding, it could represent a functional difference in the role of the CNIC and CtxIC in sound processing. One such example could be in sound localization via interaural time differences (ITD). Both human and animal research suggests that ITD in the envelope may be used in sound localization (Bernstein and Trahiotis, 2012, 2008, 1994; Klein-Hennig et al., 2011; Li et al., 2019; McFadden and Pasanen, 1976; Nuetzel and Hafter, 1976; Ono et al., 2020). The neural basis of envelope ITD encoding begins with the auditory nerve, which is well synchronized to sounds in our AM range (Joris et al., 2004). Recent research suggests that the mouse IC is sensitive to ITD in the sound envelope (Ono et al., 2020). Thus, an increase in the number of temporal modulating cells in the CNIC may indicate that the CNIC contributes more to encoding envelope ITDs than the CtxIC.

### SUPPLEMENTARY METHODS

***Lateral suppression:*** Lateral suppression was determined using simultaneous presentations of two-tone pairs (100 ms long, presented every 300 ms). In each stimulus set, one tone was presented at a fixed frequency (F1) and a simultaneously presented second tone was roved in frequency (F2; ± 0.5 to 2 octaves rel. F1; 10 steps/oct.). This was repeated for a number of F1 frequencies. Best frequency for each unit for this paradigm was approximated as the frequency in which the peak response was at F1=F2. The maximal percent suppression and the area under the curve from -0.5 to 0.5 oct. and -1 to 1 oct. was calculated. Although we calculated suppression relative to the F1 = F2 paradigm, we note that it is possible that this is an underestimate as the response to F1 alone (which is repeated on every trial) could be suppressed due to stimulus-specific adaptation (Duque and Malmierca, 2015).

***Gap detection:*** Gap detection was evaluated at a range of pure tone frequencies with gaps (0 - 100 ms, log steps) embedded 500 ms after tone onset. Stimuli were presented every 1.5 seconds. Note that because the neurons' exact BFs were only determined after offline sorting (see Discussion), the BF of each single unit for gap detection was approximated as the frequency that elicited the highest firing rate in the absence of a gap (i.e. gap length = 0 ms). The frequency corresponding to the minimum gap detection threshold was also noted. The difference in octaves from the estimated best frequency and minimum gap detection frequency was calculated. Additionally, the difference between the gap detection threshold at estimated best frequency and the minimum gap detection threshold was calculated. The gap threshold was defined as the gap length required for the firing rate during the gap to be significantly different (ANOVA) from the firing rate in an equal-length response bin immediately preceding or following the gap.

Single-unit responses were classified as onset, sustained, offset, or other based on Berger et al., 2014. The onset window was defined as the first 30 ms after stimulus onset plus the estimated onset latency (5 ms). The sustained window was defined as the 30 ms following the onset window. The offset window was defined as the first 30 ms following stimulus offset. The response in each response window was considered significant if it was greater than the spontaneous firing rate plus two standard deviations. Single units that had a significant response in only the onset window were classified as onset responders. Units with a significant response in the onset and sustained windows were considered sustained responders. Units with a significant response in only the offset window were considered offset responders. Single units that did not fit into one of these categories were classified as other.

***Responses to sinusoidally amplitude modulated tones:*** Amplitude-modulated tones (500 ms in length, AM frequencies 2 - 256 Hz, log steps, presented once every 3 seconds) were presented at a range of pure tone frequencies, based on LFP responses. The best frequency of each unit was estimated as the frequency that elicited the highest overall firing rate (see Discussion). The rate-based best modulation frequency (rBMF) and discharge synchrony-based best modulation frequency (tBMF) were calculated using previously described methods (Liang et al., 2002).
